# Neural Correlates of Attitudes and Risk Perception for Food Technology Topics

**DOI:** 10.1101/595314

**Authors:** Tyler Davis, Mark LaCour, Erin Beyer, Jessica L. Finck, Markus F. Miller

## Abstract

Food technologies provide numerous benefits to society and are extensively vetted for safety. However, many technological innovations still face high levels of skepticism from consumers. To promote development and use of food technologies, it is critical to understand the psychological and neurobiological processes associated with consumer acceptability concerns. The current study uses a neuroscience-based approach to understand consumer attitudes and perceptions of risk associated with food technologies and investigate how such attitudes impact consumer’s processing of information related to food technologies. We used functional magnetic resonance imaging (fMRI) to measure brain activation while participants processed infographics related to food technology topics. For technology topics perceived as riskier (antibiotics and hormones), activation was higher in areas of the lateral prefrontal cortex that are associated with decisional uncertainty. In contrast, technology topics that were viewed more favorably (sustainability and animal welfare) tended to activate the ventromedial prefrontal cortex, a region that processes positive affect and subjective value. Moreover, for information about hormones, the lateral PFC activation was associated with individual differences in resistance to change in risk perception. These results reveal how attitudes and risk perception relate to how the brain processes information about food technologies and how people respond to information about such technologies.

## 1. Introduction

Innovations in food technology hold great promise for improving food quality, availability, safety, and sustainability. However, like other technological innovations (Kleijnen, Lee, & Wetzelsa, 2009), new food technologies are often met with resistance in the marketplace. Consumers tend to be neophobic when it comes to foods (Pliner & Hobden, 1992), particularly foods produced with new technologies (Cox & Evans, 2008), even when expert scientists have vetted the technologies for safety (Hansen, Holm, Frewer, Robinson, & Sandøe, 2003). Such neophobia presents a barrier to development of novel technologies as well as their adoption by food producers. The high cost of investment in bringing technologies to market or deploying such technologies in food production can become financially risky when faced with the high potential for rejection by consumers (Lusk, Roosen, & Bieberstein, 2014). Thus, to promote development of safe, effective technologies for improving food sustainability and safety, it is critical to understand the psychological and neurobiological factors associated with consumer attitudes and risk perception for new technologies.

Consumer research on food technologies has uncovered several factors that influence acceptance and resistance to innovation, as well as people’s willingness-to-pay for such innovations in the marketplace (Lusk et al., 2014). At a proximal level, people’s attitudes and perceptions of risk play a critical role in influencing acceptance of new food technologies. Compared to organic or “natural” foods, foods produced with technologies are often viewed as having fewer benefits and higher risks (Frewer, Shepherd, & Sparks, 1994; Saba, Rosati, & Vassallo, 2000). Thus, consumers are often willing to pay more to avoid technologies such as genetically modified organisms (GMOs), hormones, or antibiotics than they are to purchase food created with them (Lusk et al., 2015; Lusk et al., 2014; Roosen et al., 2015). The origins of consumers’ less positive attitudes and heightened risk perception to food technologies remain elusive, but factors that are known to play a role are the perceived naturalness of foods or a technology (Tenbült, de Vries, Dreezens, & Martijn, 2005; Siegrist, 2008), tendencies to overweight low probability risks (Slovic, 2016), perceived controllability of exposure (Siegrist, Stampfli, Kastenholz, & Keller, 2008), familiarity (Tenbült, de Vries, Dreezens, & Martijn, 2008), trust in science (Lang & Hallman, 2005; Siegrist, Cousin, Kastenholz, & Wiek, 2007), and personal ethical beliefs (Sparks, Shepherd, & Frewer, 1995).

More recently, research on food technology has begun to turn to neuroscience techniques to understand the cognitive mechanisms related to consumer acceptance of food technology at a finer-grained level (Bruce et al., 2014; Lusk et al., 2015). Neuroscience research on food technologies follows the recent focus in behavioral economics on “neuroeconomic” approaches that have proven useful for understanding a number of basic economic phenomena, such as loss-aversion (Rick, 2011), temporal discounting (Kable & Glimcher, 2007; Ballard & Knutson, 2009), and how uncertainty influences preferences and perceptions of value (Platt & Huettel, 2008; Rushworth & Behrens, 2008). Multiple brain systems track basic behavioral economic constructs like value and risk, including “hot” affective systems that tend to be governed by more automatic, affective information, and “cool” systems that promote more deliberate, rational decision making (Bechara, Damasio, & Damasio, 2000; Goel & Dolan, 2003; McClure, Laibson, Loewenstein, & Cohen, 2004).

Much of neuroeconomic research has focused on the prefrontal cortex (PFC) as critical for value-based decision making processes. The prefrontal cortex is part of the frontal lobe and governs much of our higher-level cognitive functions such as planning and reasoning, as well as numerous social and affective processes. In terms of its role in value-based decision making, there are key differences within the prefrontal cortex between the lateral PFC (on the sides of the brain) and the medial PFC (the middle part of the brain, behind the forehead and above the eyes). The lateral PFC tends to track measures of economic risk and uncertainty, which are related to variability in possible outcomes associated with a decision (Kahnt, Heinzle, Park, & Haynes, 2011; Schonberg, Fox, & Poldrack, 2011). It typically activates more when a variety of different outcomes are possible and less when an outcome is completely predictable. The lateral PFC also tends to track individual differences in risk aversion, such as people’s tendency to overvalue sure bets (Christopoulos, Tolber, Bossaerts, Dolan, & Schultz, 2009). On a mechanistic level, the lateral PFC is thought to be involved in resolving conflicting information (e.g., outcome variability) in cases of higher uncertainty, due to its more general role as a center for cognitive control (Koechlin, Ody, Kouneiher, 2003; Badre & D’Esposito, 2009).

In contrast to the lateral PFC, the medial PFC, particularly the ventral part of it (the underside; ventromedial PFC), tends to track the overall subjective value of a decision (Levy & Glimcher, 2012; Bartra, McGuire, & Kable, 2013). When a decision has high subjective value, this brain region tends to activate more, and when there is low subjective value, it activates less. The medial PFC tracks individual differences in preference for immediate reward (Kable & Glimcher, 2007) and sensitivity to gains versus losses (Tom, Fox, Trepel, & Poldrack, 2007). Whereas the lateral PFC is often depicted as a “cold,” rational part of the brain, the medial part is often depicted as part of the “hot,” affective part of the brain due to its sensitivity to immediate reward (McClure et al., 2004; Krain, Wilson, Arbuckle, Castellanos, & Milham, 2006). However, both are sensitive to cognitive and affective information. The lateral PFC is involved in basic regulation of cravings and emotional responses (Delgado, Gillis, & Phelps, 2008; Kober et al., 2010; Kahathuduwa, Boyd, Davis, O’Boyle, & Binks, 2016; Kahathuduwa et al., 2018). Likewise, the ventromedial PFC integrates cognitive and more visceral affective information when computing a subjective value (Roy, Shohamy, & Wager, 2012). For example, the ventromedial PFC tracks people’s tendency to rate wines as more subjectively valuable when they are associated with a higher price tag than when the equivalent wine is low priced (“price placebo effect”; Plassmann, O’Doherty, Shiv, & Rangel 2008; Plassmann & Weber, 2015).

Following general neuroeconomic research, fMRI research on the neurobiological underpinnings of food technology choice has focused on how the PFC engages when making willingness-to-pay judgments. Consistent with the role of the lateral PFC in resolving uncertainty and conflict, lateral PFC has been found to be more active when participants are considering multi-attribute food choices that require them to consider both price and technology attributes (e.g., the price of milk and that it was produced from cloned cows; Lusk et al., 2015), and when participants opt for a higher-priced organic food to avoid a technology (Linder et al., 2010; Lusk et al., 2015). Likewise, activation in the lateral PFC tends to track relevant individual differences that contribute to price-technology conflicts, such as trait-level food technology neophobia (Bruce et al., 2014) and how much people adjust their willingness-to-pay judgments based on advertising (McFadden et al., 2015). In contrast, the ventromedial PFC has been found to be more associated with people’s final willingness-to-pay decisions in price-technology trade-offs (Crespi et al., 2015), consistent with its role in computing subjective value. In some cases, such signals from the PFC have been shown to contribute above and beyond many well-established demographic and individual difference contributors to willingness-to-pay, leading to the conclusion that functional neuroimaging provides incremental value in predicting and understanding consumer choices regarding food technology tradeoffs (Lusk et al., 2015).

With the relatively limited neuroscience research on how people perceive food technologies, there are many open questions regarding how the brain processes information about food technology and how these processes influence potential consumer behavior. Given the critical role that information and education about technologies plays in consumer acceptance, one important question is how the brain processes information about food technologies and how this processing relates to perceived risk and consumer attitudes. Perceived risk and attitudes are key pre-decisional factors that influence not only how consumers will behave when given a choice between foods produced with new technologies or more “natural” foods, but also how receptive they will be to new information (Hart et al., 2009; Sears & Freedman, 1967; Smith, Fabrigar, & Norris, 2008).

In the present study, we scanned individuals while they viewed infographics describing six food technology topics: hormones, antibiotics, vaccines, GMOs, animal welfare technologies, and sustainability technologies (for task design see Figure 1; Supplemental Figures 1-6). The infographics were selected from materials designed for marketing purposes and are publicly available on Merck Animal Health’s website prior to the study (https://www.merck-animal-health-usa.com/about/us/veterinary-consumer-affairs/animal-welfare). Thus, the infographics used are highly ecologically valid in the sense that they are from actual informational campaigns aimed at influencing consumer acceptance. Based on an online norming study (described below), we expected attitudes and risk perception to vary across the food technology topics such that hormones and antibiotics would be perceived as most risky and associated with less positive attitudes, GMOs and vaccines as intermediate, and animal welfare and sustainability technologies as least risky and associated with the most positive attitudes.

**Figure 1.**
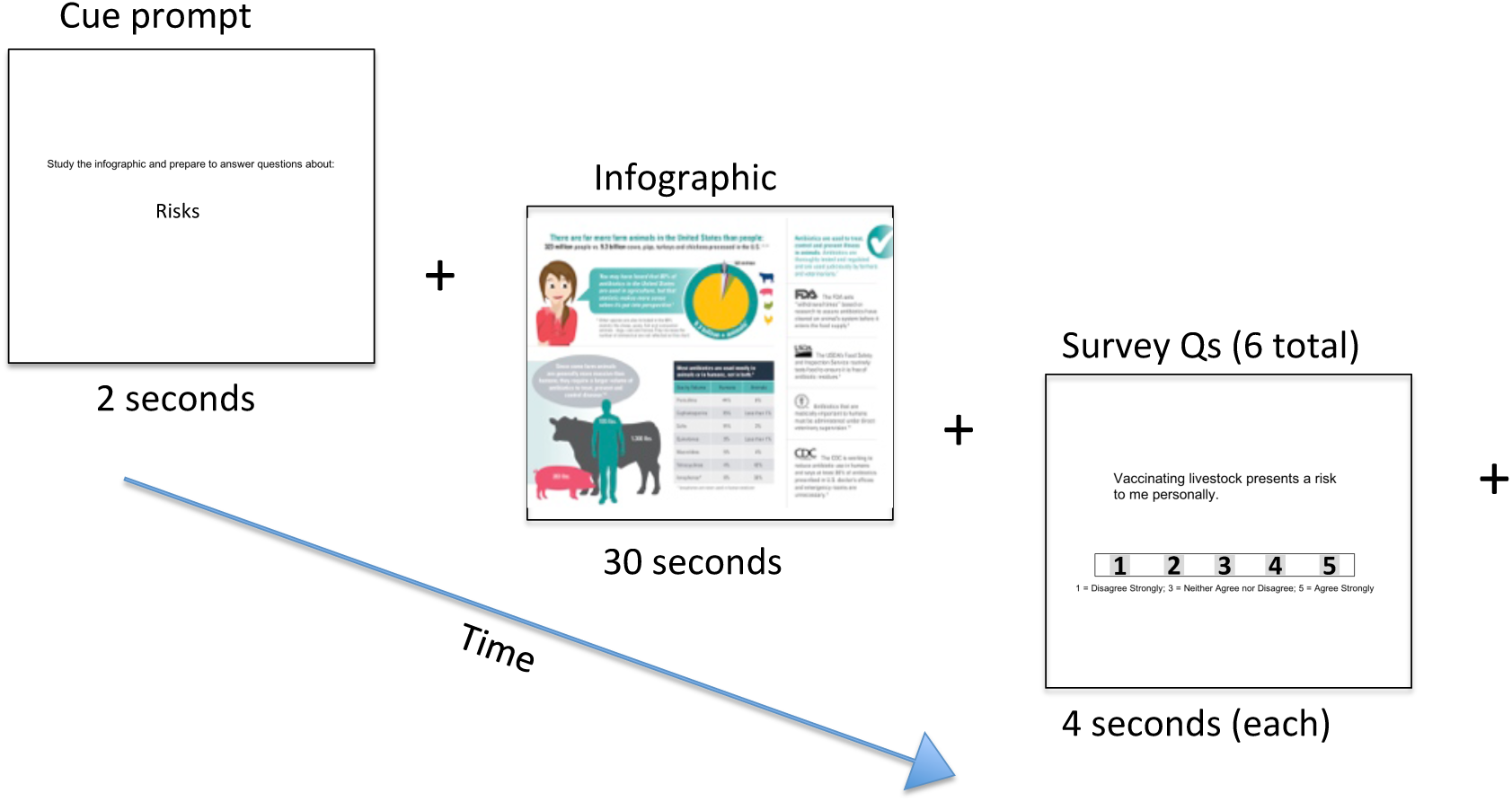
An example of the infographics presentation phase for the fMRI task. One cycle of the task would give participants a cue for what to consider in the following infographic (risks, attitudes, benefits). Then they completed 6 survey questions before the next cycle began. The +’s indicate random fixation that was interspersed to allow the fMRI signal to be separated between different task components.

In terms of the brain, we hypothesized that the same basic lateral and medial PFC regions that are involved with processing decisional information in willingness-to-pay judgments would be involved in processing the food technology infographics and that such processing would be associated with subsequent risk perception and attitudes. To this end, we hypothesized that lateral PFC activation would track the perceived riskiness of the technologies and would be associated with less positive attitudes for the technologies. Specifically, we expected that the higher perceived risk and less positive attitude technologies would be associated with more lateral PFC activation. This prediction is based on the fact that lateral PFC often tracks uncertainty and risk aversion in choice contexts (Schonberg et al., 2011). Likewise, we hypothesized that ventromedial PFC activation would track positive attitudes such that the technologies with the most positive attitudes would activate the ventromedial PFC more. This prediction is based on the fact that ventromedial PFC often tracks liking and positive value, even outside of choice contexts (Roy et al., 2012).

In addition to our primary goals, which are to examine how risk perception and positive attitudes relate to brain activation during infographic processing, we also examine how individual differences in risk perception and attitude change (from pre-to post-infographic) are associated with individual differences in brain activation. These individual difference effects can augment the primary associations by answering whether, for example, the hypothesized risk-related processing in lateral PFC is generally associated with improving risk perception and attitudes or if it is associated with resisting such change.

## 2. Materials and Methods

### 2.1 Participants

Three hundred and ninety four participants completed an online version of the study (online norming study) to provide normative data for the neuroimaging study. These participants were recruited through Amazon’s Mechanical Turk and paid $4 for participation. Fifty-three participants completed the neuroimaging study. Neuroimaging participants were paid $25 and given an extra $5 for a willingness-to-pay auction that was administered after all of the infographic fMRI scans were completed. All protocols were approved by the Human Rights Protection Program (HRPP) of Texas Tech University (Proposal number: IRB2018-730; Approved: 9/6/2018).

### 2.2 Behavioral Protocols

#### 2.2.1 Online Norming Study

The online norming study (Figure 2) consisted of a demographic section, attitudes and risk perception survey sections for each of the following technology/animal welfare topics: antibiotics, vaccines, hormones, GMOs, sustainability, and animal welfare, followed by a set of non-technology specific standardized questionnaires: Food Technology Neophobia, Trust-in-Science, and Chemical Reasoning.

**Figure 2.**
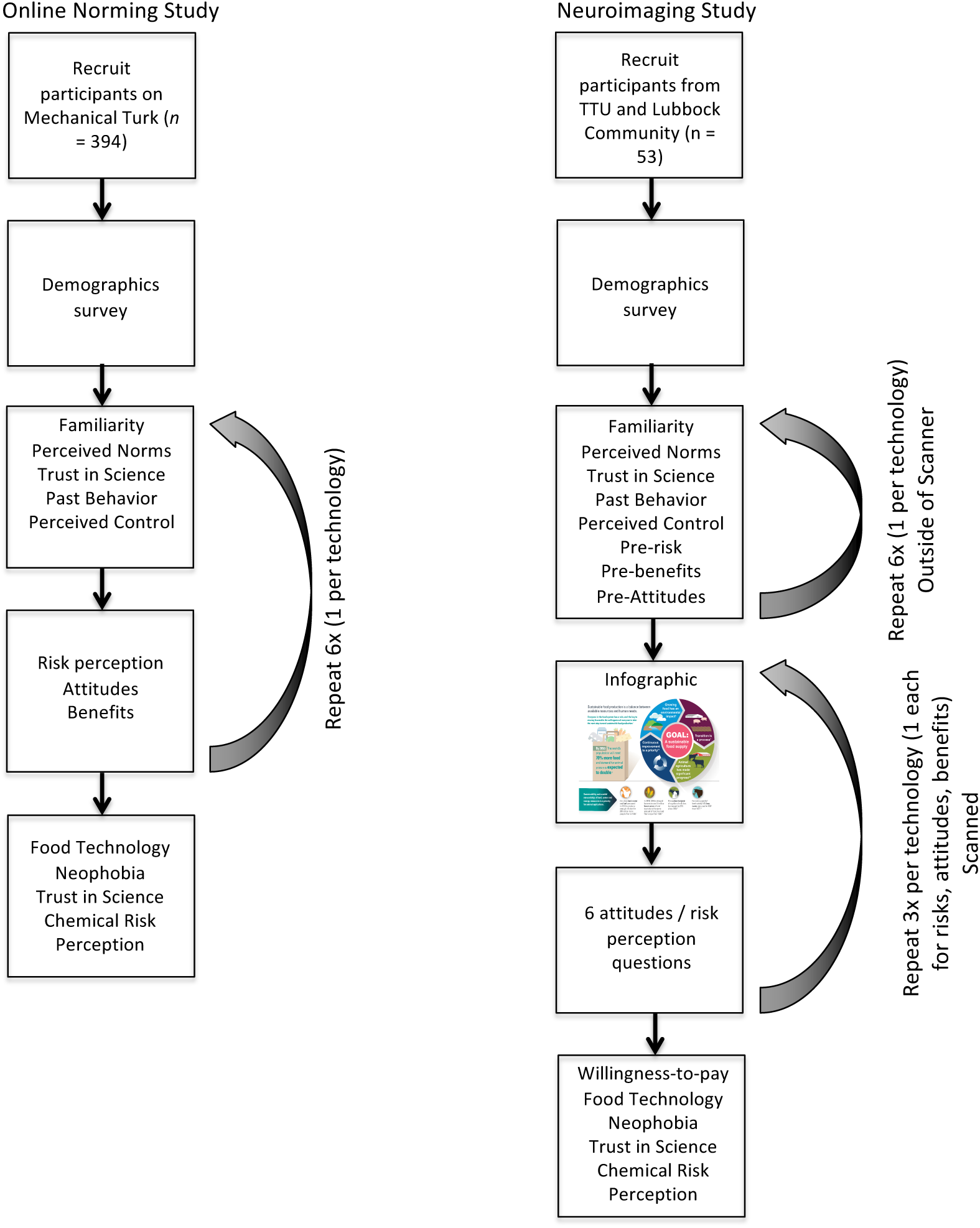
A depiction of the study flow for the online norming and neuroimaging studies.

The demographics section included questions asking people to report their age, sex, ethnicity, first language, zip code (or country) in which they spent majority of childhood, highest level of education completed, political orientation, organic buying behavior, whether they eat meat, and whether they subscribe to any dietary philosophies (vegan, vegetarian, etc.; see Table 1).

**Table 1.**
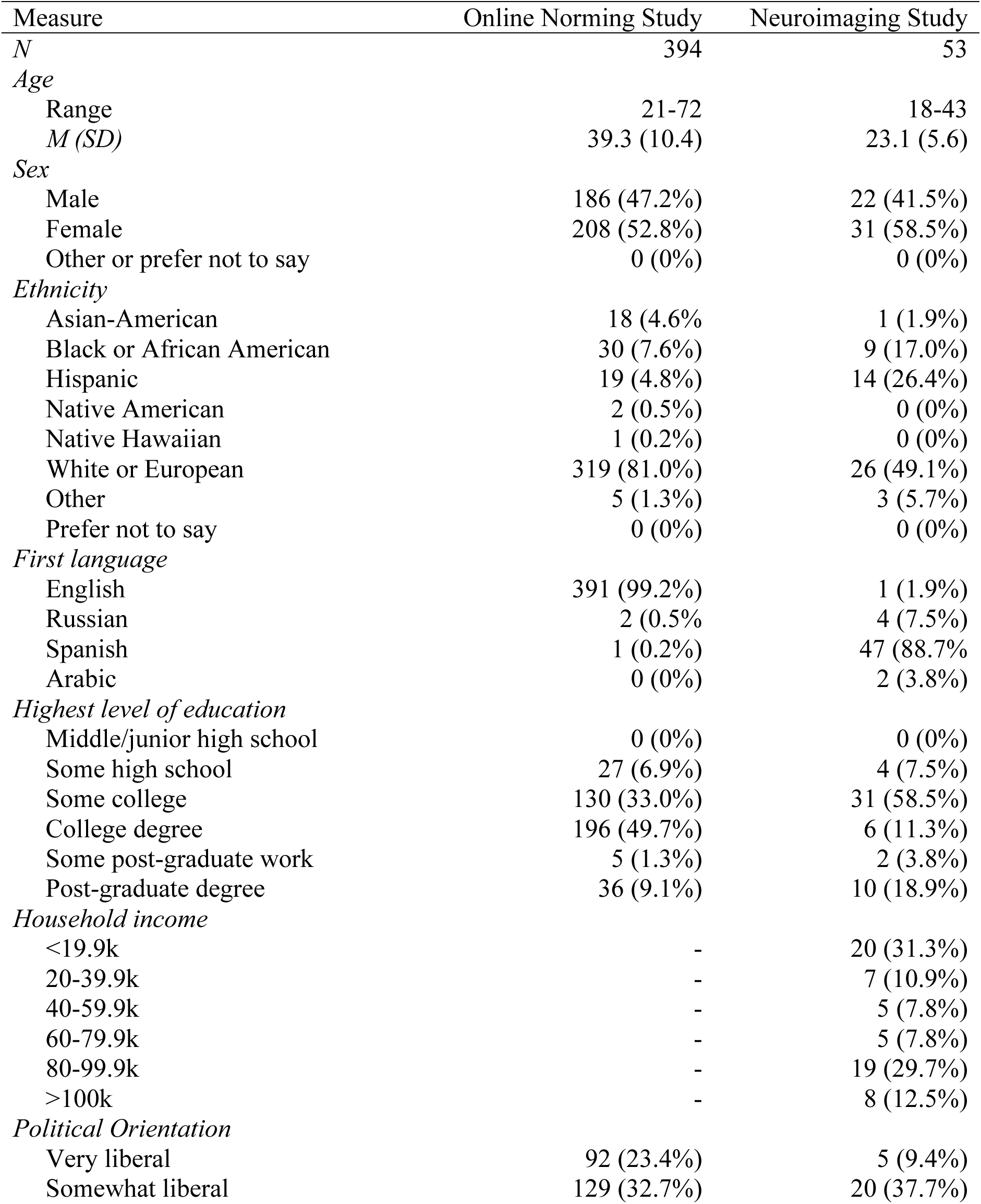
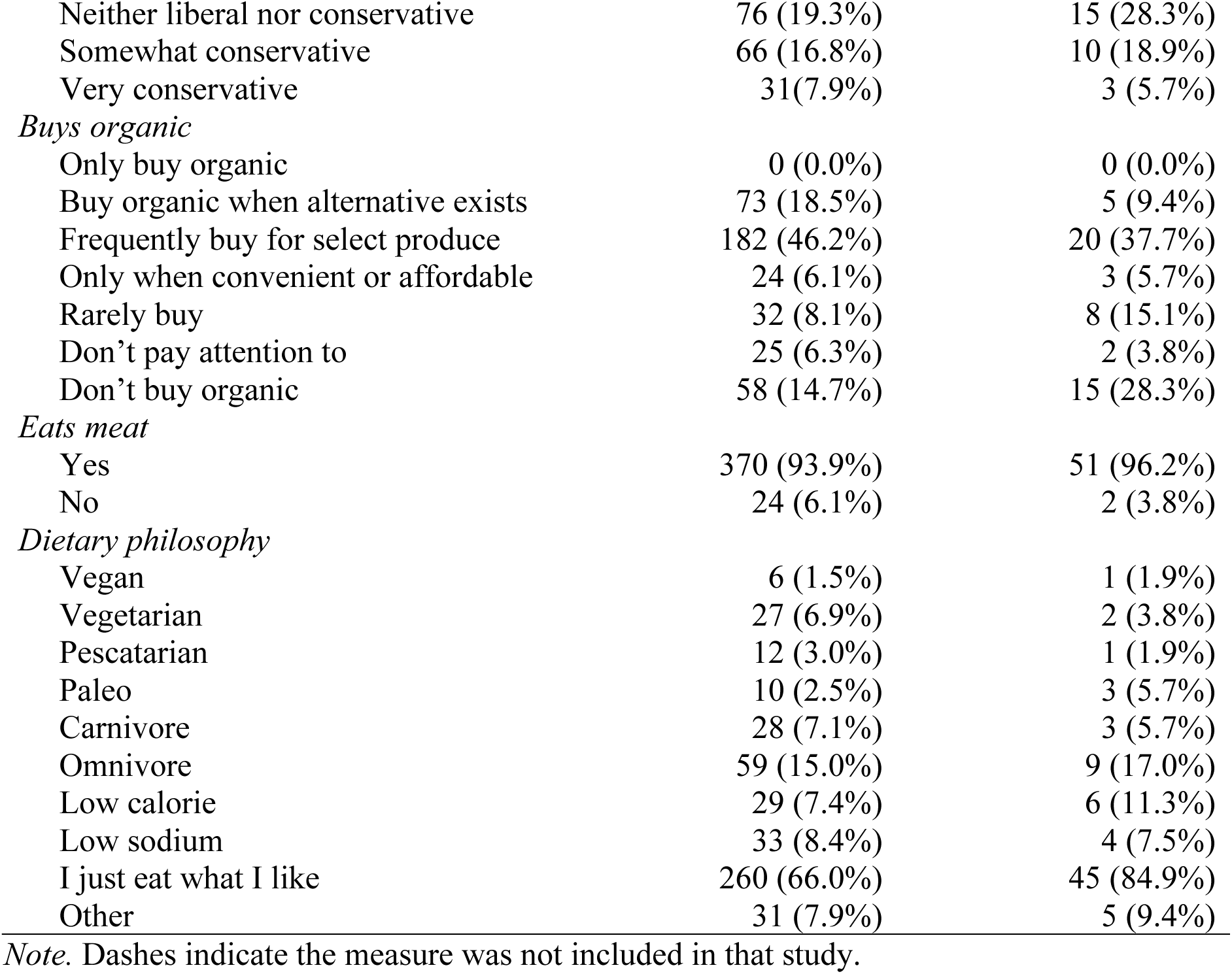
Demographics and Baseline Food Preferences

| Measure | Online Norming Study | Neuroimaging Study |
| --- | --- | --- |
| *N* | 394 | 53 |
| *Age* |  |  |
| Range | 21-72 | 18-43 |
| *M (SD)* | 39.3 (10.4) | 23.1 (5.6) |
| *Sex* |  |  |
| Male | 186 (47.2%) | 22 (41.5%) |
| Female | 208 (52.8%) | 31 (58.5%) |
| Other or prefer not to say | 0 (0%) | 0 (0%) |
| *Ethnicity* |  |  |
| Asian-American | 18 (4.6%) | 1 (1.9%) |
| Black or African American | 30 (7.6%) | 9 (17.0%) |
| Hispanic | 19 (4.8%) | 14 (26.4%) |
| Native American | 2 (0.5%) | 0 (0%) |
| Native Hawaiian | 1 (0.2%) | 0 (0%) |
| White or European | 319 (81.0%) | 26 (49.1%) |
| Other | 5 (1.3%) | 3 (5.7%) |
| Prefer not to say | 0 (0%) | 0 (0%) |
| *First language* |  |  |
| English | 391 (99.2%) | 1 (1.9%) |
| Russian | 2 (0.5%) | 4 (7.5%) |
| Spanish | 1 (0.2%) | 47 (88.7%) |
| Arabic | 0 (0%) | 2 (3.8%) |
| *Highest level of education* |  |  |
| Middle/junior high school | 0 (0%) | 0 (0%) |
| Some high school | 27 (6.9%) | 4 (7.5%) |
| Some college | 130 (33.0%) | 31 (58.5%) |
| College degree | 196 (49.7%) | 6 (11.3%) |
| Some post-graduate work | 5 (1.3%) | 2 (3.8%) |
| Post-graduate degree | 36 (9.1%) | 10 (18.9%) |
| *Household income* |  |  |
| <19.9k | - | 20 (31.3%) |
| 20-39.9k | - | 7 (10.9%) |
| 40-59.9k | - | 5 (7.8%) |
| 60-79.9k | - | 5 (7.8%) |
| 80-99.9k | - | 19 (29.7%) |
| >100k | - | 8 (12.5%) |
| *Political Orientation* |  |  |
| Very liberal | 92 (23.4%) | 5 (9.4%) |
| Somewhat liberal | 129 (32.7%) | 20 (37.7%) |
| Neither liberal nor conservative | 76 (19.3%) | 15 (28.3%) |
| Somewhat conservative | 66 (16.8%) | 10 (18.9%) |
| Very conservative | 31 (7.9%) | 3 (5.7%) |
| *Buys organic* |  |  |
| Only buy organic | 0 (0.0%) | 0 (0.0%) |
| Buy organic when alternative exists | 73 (18.5%) | 5 (9.4%) |
| Frequently buy for select produce | 182 (46.2%) | 20 (37.7%) |
| Only when convenient or affordable | 24 (6.1%) | 3 (5.7%) |
| Rarely buy | 32 (8.1%) | 8 (15.1%) |
| Don't pay attention to | 25 (6.3%) | 2 (3.8%) |
| Don't buy organic | 58 (14.7%) | 15 (28.3%) |
| *Eats meat* |  |  |
| Yes | 370 (93.9%) | 51 (96.2%) |
| No | 24 (6.1%) | 2 (3.8%) |
| *Dietary philosophy* |  |  |
| Vegan | 6 (1.5%) | 1 (1.9%) |
| Vegetarian | 27 (6.9%) | 2 (3.8%) |
| Pescatarian | 12 (3.0%) | 1 (1.9%) |
| Paleo | 10 (2.5%) | 3 (5.7%) |
| Carnivore | 28 (7.1%) | 3 (5.7%) |
| Omnivore | 59 (15.0%) | 9 (17.0%) |
| Low calorie | 29 (7.4%) | 6 (11.3%) |
| Low sodium | 33 (8.4%) | 4 (7.5%) |
| I just eat what I like | 260 (66.0%) | 45 (84.9%) |
| Other | 31 (7.9%) | 5 (9.4%) |

*Note.* Dashes indicate the measure was not included in that study.

The attitudes and risk perception survey sections went through each technology, one at a time, such that all attitudes questions were asked about a given technology prior to beginning the next. The online norming group did not see any infographics. The order of the technologies was balanced across participants using a Latin square to neutralize potential order effects. For each technology, the attitudes surveys began with a brief description of the technology, which participants were required to briefly summarize in their own words prior to moving on (to ensure participants were paying attention). Next, participants were asked two questions each on familiarity, subjective norms, trust in science (with respect to the specific technology), past behavior with the technology (consuming or avoiding food produced with the technology), and perceived control. All of these factors are known to potentially impact attitudes and risk perception (Aizen, 1991; Slovic, 2016). We used wordings that are consistent with these existing risk perception studies and the Theory of Planned Behavior (see Supplemental Table 1 for wording for all questionnaires). However, in some cases we had to adapt wordings from typical wordings to ensure that the same basic questions could be used for all of the technologies of interest. After the preliminary questions, participants answered a section on risk perception and two sections on attitudes: one on more personal attitudes and one on more general benefits. Each of the risk and attitude sections contained eight questions, two of which were designated as “pre” questions and added to the pre-surveys for the neuroimaging study, and six were designated as “post” questions and asked after the infographic exposure in the neuroimaging study. Because no infographics were shown in the online norming study, both “post” and “pre” questions were asked at the same time in the online norming study. Examples of the wording of all questions in the attitudes sections are listed in Supplemental Table 1.

After completing the technology-specific questions for each of the six technologies, participants completed standardized surveys on Food Technology Neophobia scale (Cox and Evans, 2008), a standardized Trust in Science Scale (Bak, 2001), and a set of questions from a study on chemical risk perception (Kraus, Malmfors, & Slovic, 1992) that includes factors related to dose-to-response reasoning, attitudes toward chemicals, chemical risk reduction, and miscellaneous questions on chemical risk perception.

#### 2.2.2 Neuroimaging Study

The neuroimaging study (Figure 2) included a demographic section, a pre-infographic exposure survey taken outside of the scanner, infographic exposure and attitude/risk perception surveys done inside the scanner for each technology, and a set of willingness-to-pay judgments and non-technology specific standardized questionnaires (Food Technology, Neophobia, Trust-in-Science, chemical risk perception) taken after all of the infographic exposures and technology specific attitudes/risk perception surveys.

The demographic section was the same as that used for the online norming study, except we also included a question asking for household income (Table 1). After the demographic section, the participants completed a pre-infographic survey for each technology that included all of the questions used in the norming study for familiarity, perceived norms, technology-specific trust-in-science, past behavior with the technology, and perceived control, as well as two pre-infographic questions on each of general attitudes, benefits, and risk perception.

In the infographic and attitude/risk perception section (see Figures 1 and 2), infographics depicting the six food technology topics (antibiotics, vaccines, hormones, GMOs, sustainability, and animal welfare; See Supplemental Figures 1-6 for the infographics used in the study) were presented to participants separately in different scanning runs. The order of infographics was balanced across participants using a Latin square. Within a run, participants were presented with an infographic depicting one of the technologies three times. Prior to each infographic presentation, participants were cued/primed (2 seconds) with what information they would be rating next (“attitudes”, “benefits”, and “risks”). Priming participants about the type of questions they would be answering next was done to encourage participants to actively process the infographics during the presentation blocks and to scan all of the information in the infographic across the three presentations. The cues were separated by variable duration fixation cross (2, 3, or 4s), which participants were simply told to keep their eyes on while awaiting infographics. The fixation cross provides a baseline that helps to statistically isolate the responses to the different task components. After fixation, the infographic was presented for 30s. After the infographic was presented, fixation was briefly presented (5, 6, or 7 seconds) and then six post-infographic Likert-type questions about risks or attitudinal information (attitudes or benefits) were presented for 4 seconds each. During these questions, participants keyed in their responses on a sliding scale that ranged from 1 (*strongly disagree*) to 5 (*strongly agree*) by using an MR-safe button box held in their right hand. Finally, the risk and attitudes questions were also separated by fixation, which averaged 4 seconds and was drawn from a truncated exponential distribution. A second and third infographic cue, infographic presentation, and question period would then begin until all questions were answered for an infographic.

After the primary infographic runs, participants completed a set of willingness-to-pay judgments and the same Food Technology Neophobia, Trust-in-Science, and chemical risk perception surveys as were used in the online norming study. The willingness-to-pay judgments consisted of a Becker-DeGroot-Marschack (1964) style auction. In this auction, participants were asked to bid from $1-$5 on a variety of food items produced with food technologies (see Supplemental Figure 7 for an example trial). Participants were told that one of the food items would be selected randomly at the conclusion of the auction for purchase using their $5 endowment. They were told that the cost of the item would be whatever they bid, and that the probability a specific food item would be selected was based on how much they bid, such that a $5 bid was 5x more likely to win than a $1 bid. They were also told to consider the portion for each food item to be worth $5 retail and thus the rational choice was to bid however much they were willing to pay for each item. The foods the participants evaluated were beef jerky, turkey jerky, canned chicken, chili, coffee creamer, and frosting. Participants considered these foods with each of the following technology related prompts: “Animals raised fully organic”, “Animals raised all natural”, “Animals raised with hormone implants”, “Animals previously vaccinated”, “Animals previously treated with antibiotics”, “Animals raised with genetically modified (GM) feed”, “Animals raised with animal welfare technologies”, and “Animals raised with sustainable food technologies”. Because some of the technology options do not exist for particular foods (e.g., hormone implants are only used in beef production) or are impossible to verify through labeling (type of feed), at the end of the study, participants were told that they could simply select one of whatever food item they wanted from conventional, “natural” labeled, or organic labeled versions of each of the food options. Six participants did not have willingness-to-pay judgments properly recorded and were excluded from that analysis.

### 2.3 Image Acquisition

Imaging data were acquired at Texas Tech Neuroimaging Institute on a 3T Siemens Skyra using a 20-channel head-coil. Functional images were acquired in the axial plane using the Siemen’s product EPI sequence: TR = 2.09 s; TE = 25 ms; θ = 70°; FoV= 192 x 192 mm; matrix = 64 x 64; number of slices = 41, slice thickness = 2.5 mm; 0.5 mm gap. Slices were tilted to reduce orbitofrontal dropout (Deichmann, Gottfried, Hutton, & Turner, 2003). In addition, a high-resolution anatomical image was collected for each of the participants using an MPRAGE sequence acquired in the sagittal plane: TR = 1.9 s; TE = 2.49 ms; *θ* = 9°; FoV = 240 x 240 mm; matrix = 256 x 256 mm; slice thickness = 0.9 mm, slices = 192.

#### 2.4.1 Behavioral Data Analysis

Participant averages were calculated for each of the behavioral measures by averaging across the respective scale items for each construct (e.g., averaging across all post-infographic attitudes questions for hormones to get a single hormone attitude average for each participant). Group-level averages and standard deviations were then calculated for each measure by averaging across participant means. Benefits and personal attitudes were combined a priori into a single positive attitudes measure for all analysis (behavioral and neuroimaging) because there were no *a priori* differences predicted between benefits and personal attitudes in terms of brain processing or behavior, and because the combination of personal attitudes and benefits only increased reliability (see Supplemental Table 2 for Cronbach’s coefficient alpha for all risk and attitude measures).

Behavioral data from both studies were analyzed using linear mixed-effects models implemented in R’s nlme package (Pinheiro and Bates, 2006). The overall analysis strategy was to assess whether participants’ mean risk perception, attitudes, and willingness-to-pay varied significantly across the food technologies. Follow-up t-tests were used to test specific planned contrasts for how these measures varied across the technologies. Specifically, for risk perception and attitudes, we tested whether hormones and antibiotics were associated with higher risk perception and less positive attitudes compared to (1) GMOs and vaccines and (2) animal welfare and sustainability technologies. We also tested (3) whether GMOs and vaccines were associated with higher risk perception and lower positive attitudes than sustainability and animal welfare technologies. For the willingness-to-pay judgments, we tested whether participants were more willing to pay for organic and natural foods relative to (1) animals raised/treated with antibiotics and hormones, (2) animals raised/treated with GM feed and vaccines, and (3) animals raised with sustainability and animal welfare technologies.

To test whether attitudes or risk perception changed from pre-to post-infographic in the neuroimaging study, we adopted a standardization procedure using the online norming data. Z-scored risk perception (*Zrisk*) and attitude (*Zatt*) variables were calculated for each participant *j* and technology *t* by:

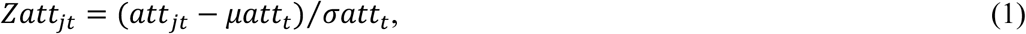

where *att_jt_* is neuroimaging participant *j*’s mean positive attitude rating for technology *t*, att_t_ is the mean positive attitudes rating for technology *t* from the online norming study, and *att_t_* is the standard deviation of positive attitude ratings for technology *t* from the online norming study. Four Z values were calculated for each technology and neuroimaging participant: one for each pre- and post-infographic scale for risk perception and attitudes. The purpose of this standardization procedure was to neutralize mean differences between the pre- and post-attitudes questionnaires that were simply due to differences in the pre and post question means. By removing the online norming study’s means, any remaining differences between pre- and post-infographic should be due to changes in risk perception or attitudes from viewing the infographics. The differences between pre- and post-infographic Z’s (post-pre) were tested using separate t-tests for each technology. The Z difference (post-pre) measures are also used in the fMRI analysis testing for individual differences in risk perception and attitude change (see *2.4.2 fMRI Data Analysis*).

#### 2.4.2 fMRI Data Analysis

FMRI analysis was conducted using a standard three-level general linear model (GLM) implemented in FEAT (FMRI Expert Analysis Tool) Version 6.00, part of FSL (FMRIB’s Software Library, www.fmrib.ox.ac.uk/fsl). Data were first pre-processed using rigid-body motion correction and skull stripping. The first-level model included explanatory variables (EVs) for cue presentation, infographic presentation (separate EVs for risk and attitudes prompts), stimulus presentation for risk and attitudes questions, and participants’ ratings as a parametric modulator of the rating questions. First-level models also included nuisance regressors for motion (6 motion parameters and their temporal derivatives) and regressors to “scrub” (censor) volumes with framewise displacement > .9mm (Siegel et al., 2014). The first level model also included temporal derivatives for each task variable, high-pass filtering, spatial smoothing (6mm FWHM), two-stage registration to high resolution anatomical and standard space, and pre-whitening to account for temporal autocorrelation. Second level models included EVs for each infographic type (antibiotics, vaccines, hormones, GMOs, sustainability, and animal welfare) and included one contrast weighting each infographic type by its average risk rating, and one contrast weighting each infographic by its average attitude rating (ratings were z-scored to ensure contrast weights summed to zero). These contrasts estimate, for each participant, which brain regions correlate with between-infographic differences in risk perception and attitudes, respectively. Third-level models tested the average group-level effect of the second-level contrasts, treating participant as a random effect for population inference. Additional third-level models were used to test whether brain activation for any of the infographics was associated with change in attitudes or risk perception from pre-to post-infographic by correlating activation with participants’ post – pre infographic Z scores (see *2.4.1 Behavioral Data Analysis*).

The primary (cluster-forming) threshold for statistical maps was set at *z* = 3.1 (*p* = .001, one-tailed), and corrected for multiple comparisons at p < .05 using a cluster-extent threshold (number of contiguous voxels exceeding the z = 3.1 cutoff) based on Gaussian Random Field Theory. Our hypothesized regions-of-interest (ROIs) were the lateral prefrontal cortex and ventromedial prefrontal cortex; however, all statistical analyses were conducted and corrected for multiple comparisons at the whole-brain level to allow for the possibility that additional regions would track infographic-level differences in risk perception and/or positive attitudes.

## 3. Results

### 3.1 Behavioral Results

#### 3.1.1 Online Norming Study

Our primary question for the online norming study was whether the technologies differed in terms of mean positive attitudes and risk perception (Table 2). Using linear mixed effects models, we found significant variability in the technology means across each of the scale measures (Table 3). Note that the pre- and post-scales are separated to align with the neuroimaging study, but because no infographics were presented in the online norming study, all measures were taken at the same time. Planned contrasts revealed that hormones and antibiotics were perceived as significantly riskier and associated with significantly less positive attitudes compared to (1) vaccines and GMOs and (2) sustainability and animal welfare technologies.

**Table 2.**
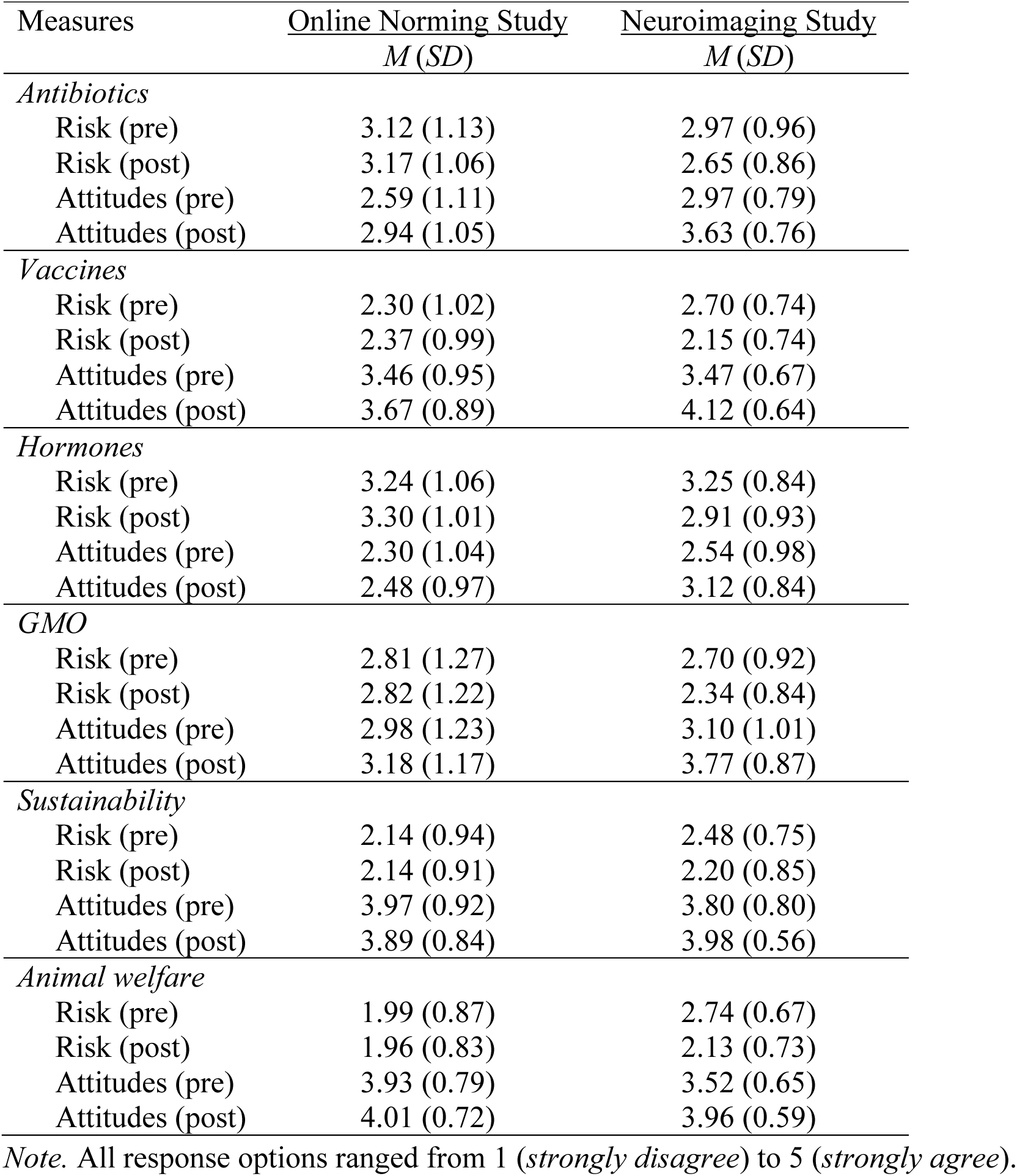
Mean and Standard Deviations for Technology-Specific Survey Measures Across studies

**Table 3.**
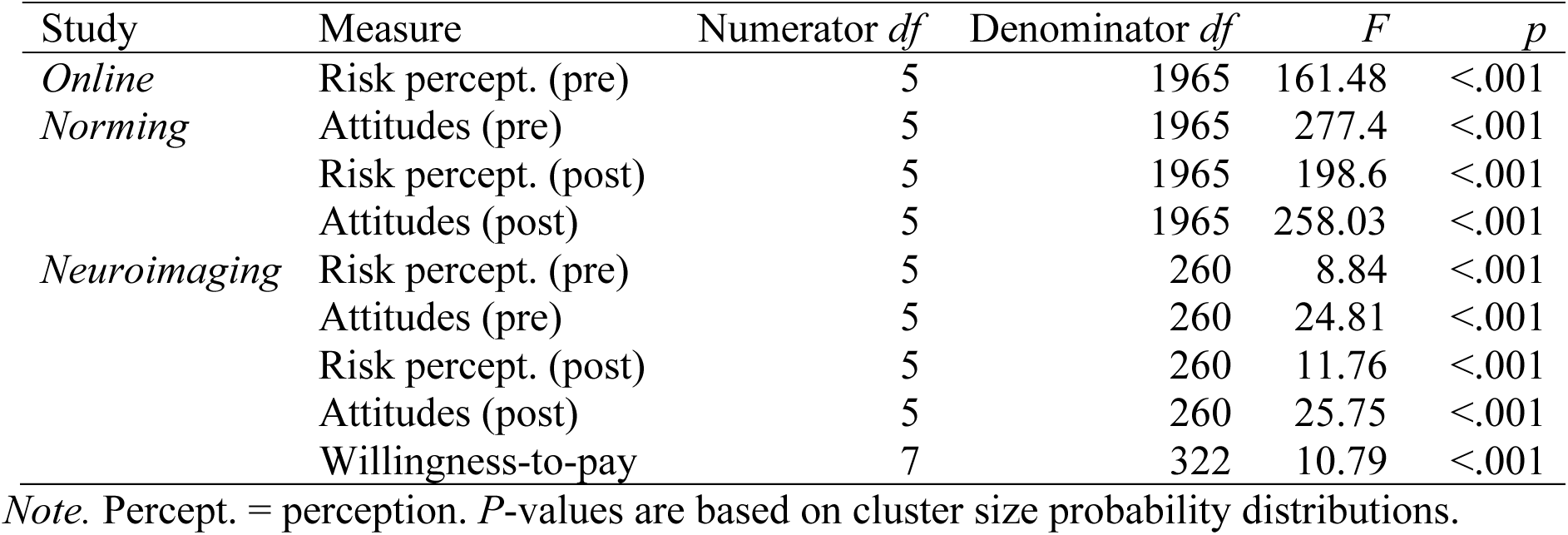
Multilevel Model Summaries for Online Norming and fMRI Studies

Vaccines and GMOs were perceived as significantly riskier and associated with significantly less positive attitudes compared to sustainability and animal welfare technologies (Table 4). Means and standard deviations for other survey measures are listed in Supplemental Tables 3 & 4.

**Table 4.**
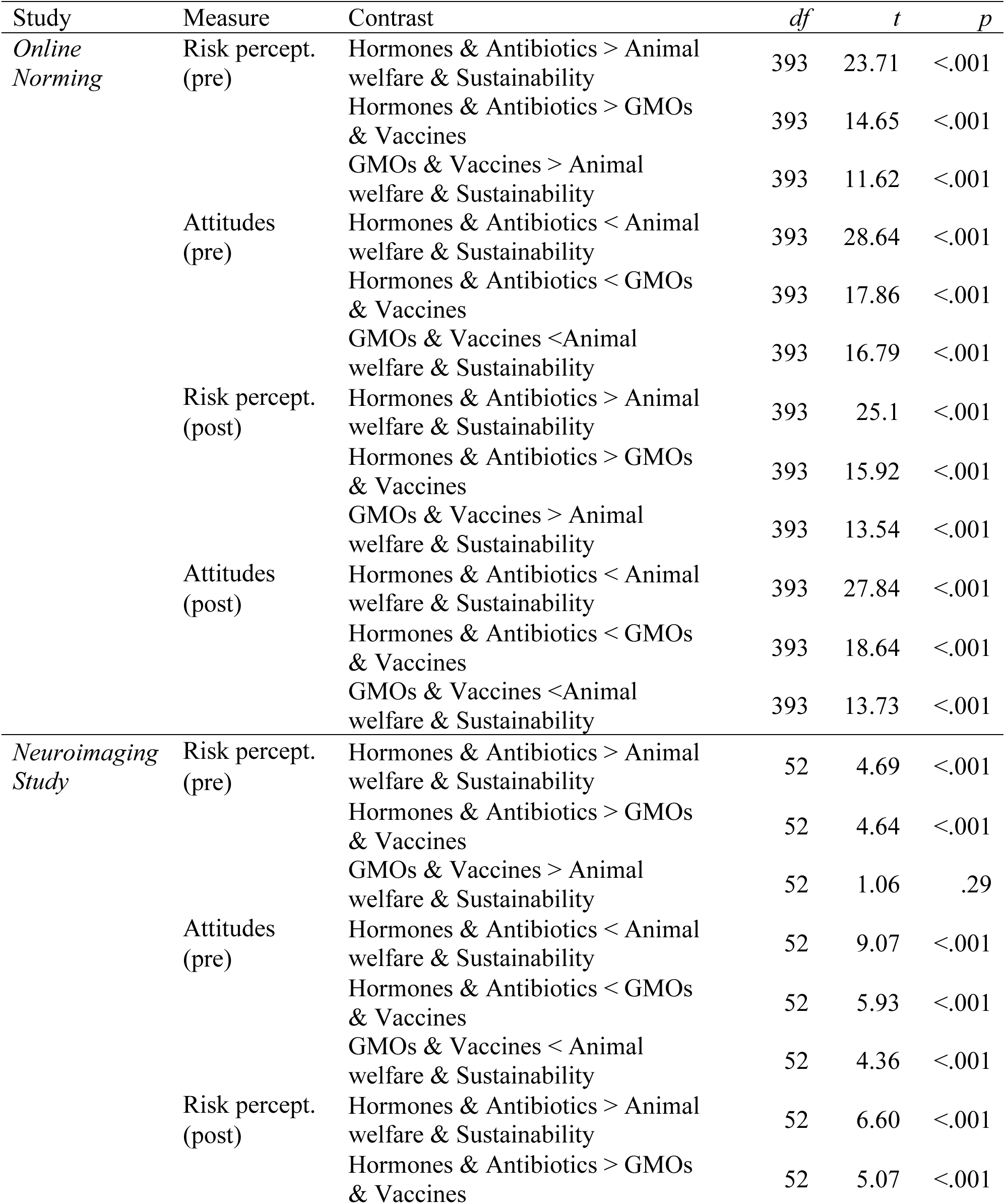
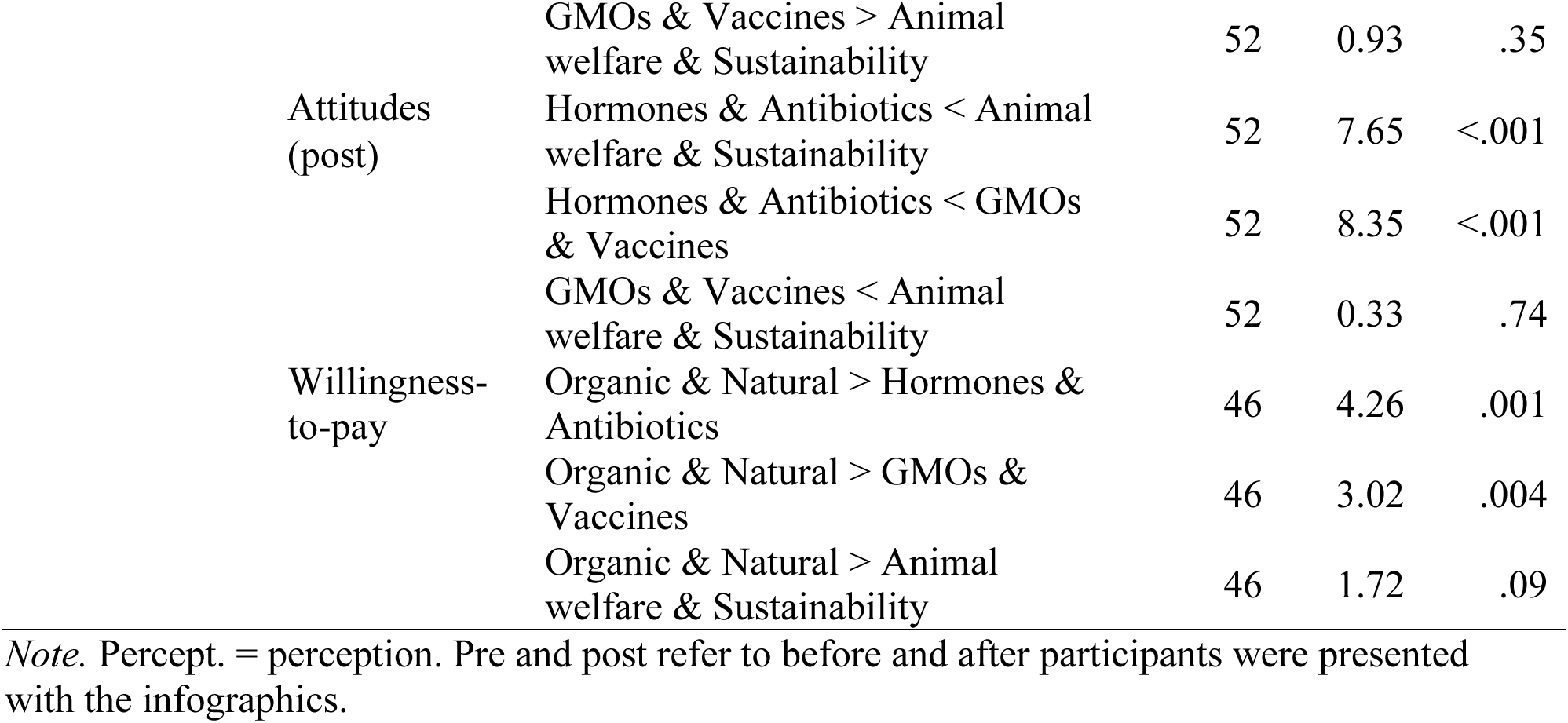
Test Statistics for Planned Contrasts for the Online Norming Study and the fMRI Study

#### 3.1.2 Neuroimaging Study

The behavioral results of the neuroimaging study were similar to the online norming study. There was significant variability across the technologies in all risk perception and attitudes measures (pre- and post-infographic; Tables 2-3; Figure 3). Like the online norming study, hormones and antibiotics were perceived as significantly riskier and were associated with less positive attitudes compared to (1) vaccines and GMOs and (2) sustainability and animal welfare technologies. However, vaccines and GMOs did not significantly differ from sustainability and animal welfare technologies for any measures except for the pre-infographic attitudes measure where they were associated with significantly less positive attitudes (Table 4). Comparing post-infographic to-pre-attitudes using Z difference scores (post – pre) that were normalized with respect to the online norming study’s scale means and standard deviations revealed that risk perception decreased and positive attitudes increased for all technologies after viewing the infographics (Table 5). Means and standard deviations for other survey measures are listed in Supplemental Table 3.

**Figure 3.**
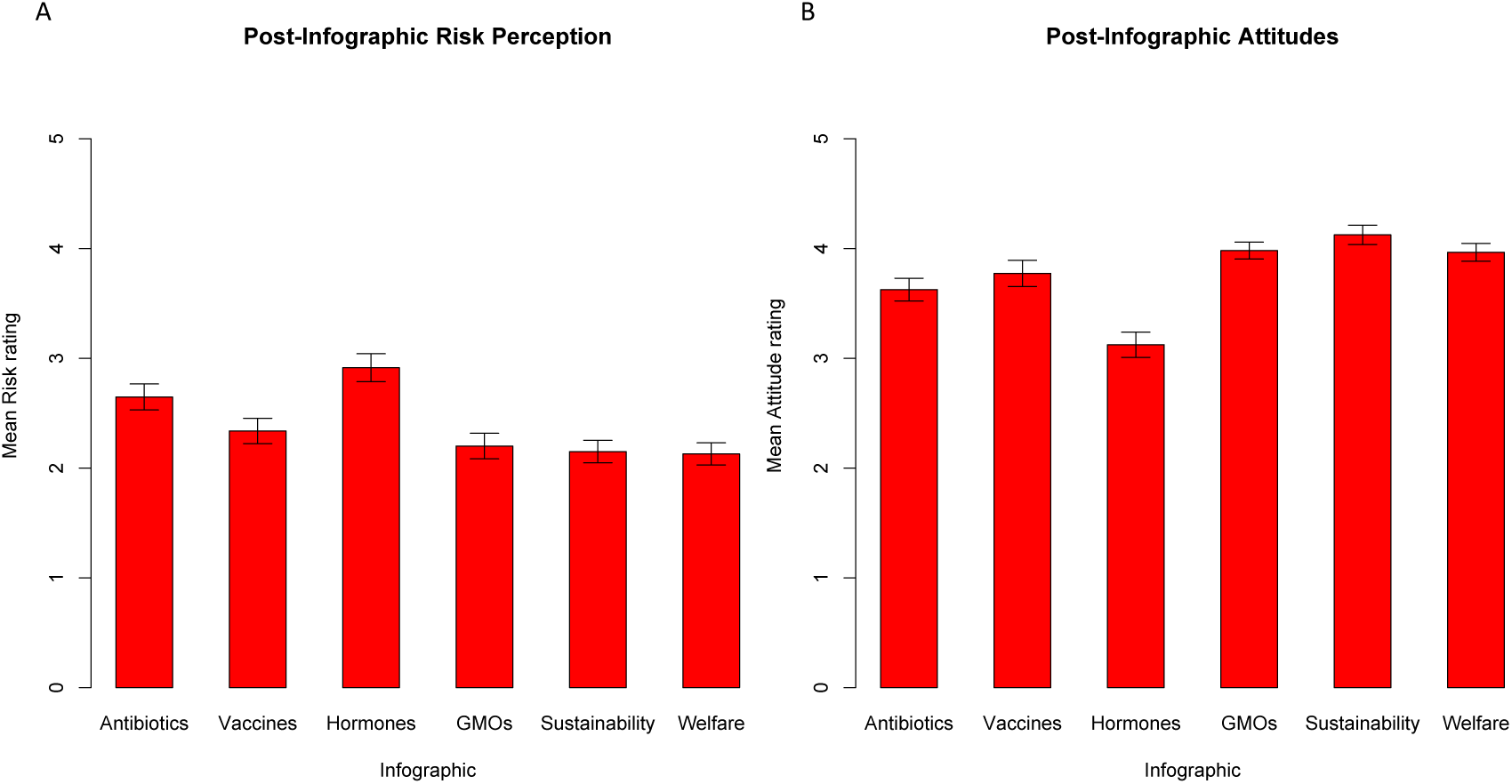
Mean post-infographics behavioral ratings for risk (A) and attitudes/benefits (B). The ratings were obtained during the scanning blocks after the infographic presentations. The error bars indicate +/- 1 standard error of the mean. The y-axis ranges from 1-5 which includes the full range of each rating scale.

**Table 5.**
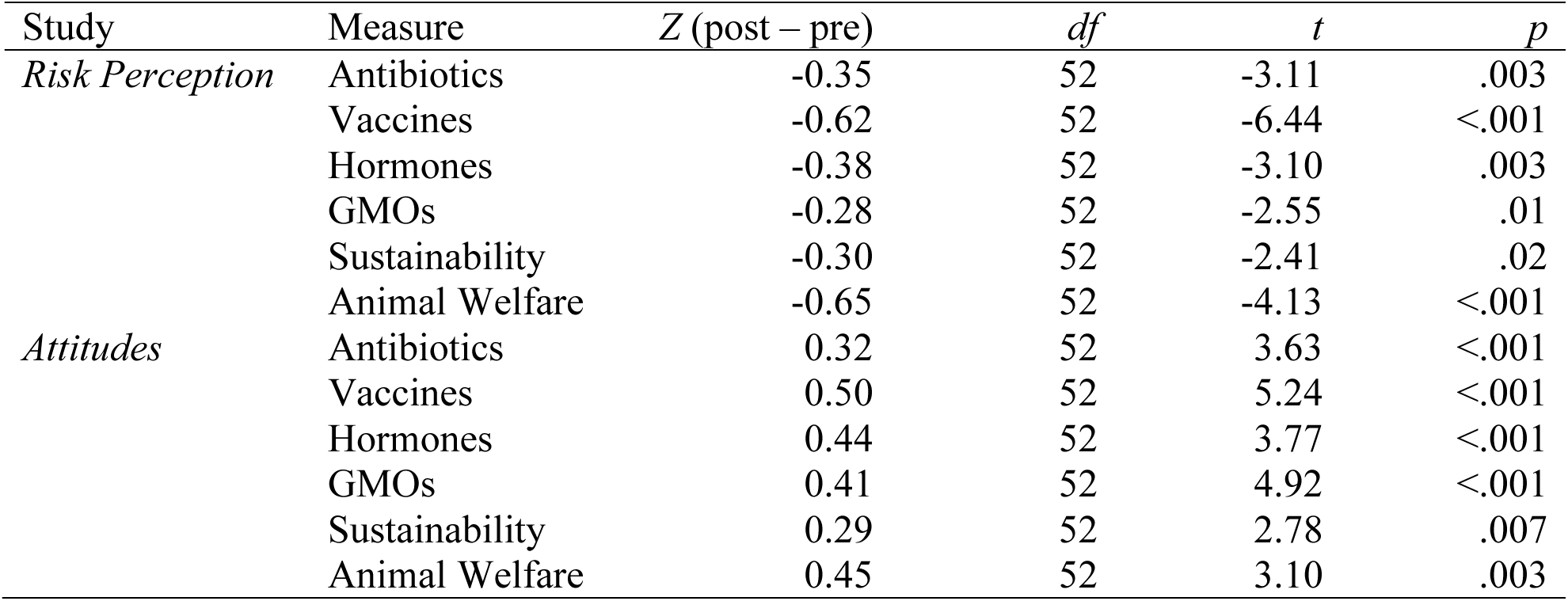
Post – Pre-Infographic Differences in Risk and Attitudes

Analysis of willingness-to-pay judgments also revealed significant variability across the technologies (Table 2; Figure 4), with willingness-to-pay for organic and natural labeling exceeding (1) hormones and antibiotics, (2) vaccines and GMOs, and (3) sustainability and animal welfare technologies (Tables 4 & 6).

**Figure 4.**
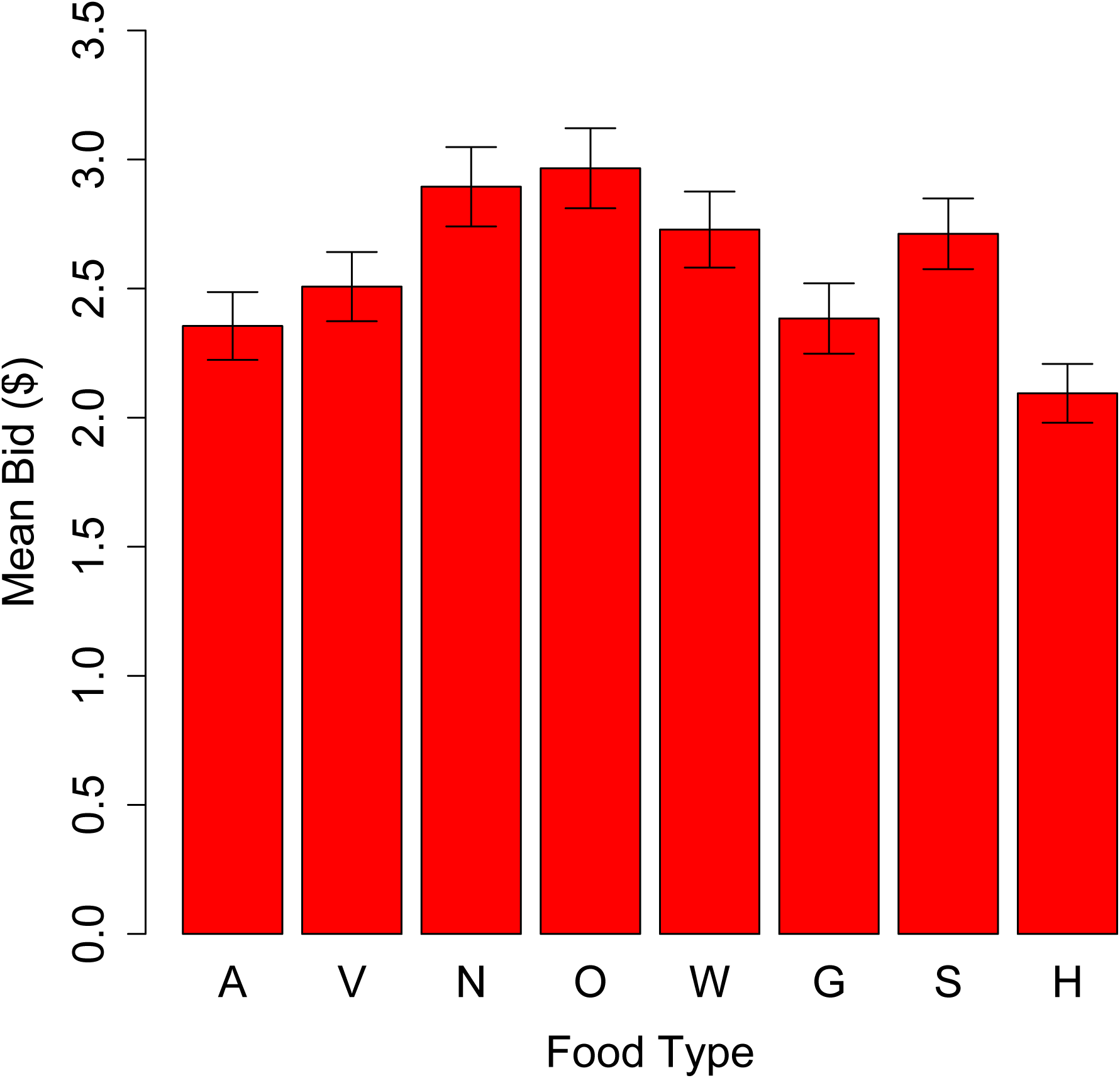
Mean willingness-to-pay judgments from the Becker-DeGroot-Marschack (1964) style auction in the neuroimaging study. A = Antibiotics, V = Vaccines, N = All natural, O = Organic, W = Animal welfare, G = GMO, S = Sustainability, H = Hormones. The error bars indicate +/- 1 standard error of the mean. The y-axis ranges from $0 – $3.5 due to the small range of averages, but bids were allowed up to $5.

**Table 6.** Willingness-to-Pay Judgments from Becker-DeGroot-Maschak Style Auction

| Target | *M (SD)* |
| --- | --- |
| Animals previously treated with antibiotics | 2.36 (0.96) |
| Animals previously vaccinated | 2.51 (0.98) |
| Animals raised all natural | 2.89 (1.12) |
| Animals raised fully organic | 2.97 (1.13) |
| Animals raised with animal welfare technologies | 2.73 (1.08) |
| Animals raised with genetically modified feed | 2.38 (0.99) |
| Animals raised with sustainable food technologies | 2.71 (1.00) |
| Animals raised with hormone implants | 2.09 (0.83) |

*Note.* All payment judgments were in US dollars. The possible range was \$1-\$5 for each item.

### Between-infographic differences in activation

#### 3.2.1 Activation related to risk processing

Four clusters of activation were positively associated with mean risk ratings across infographics (greater activation for higher risk; Figure 5; Table 7): one spanning the left dorsolateral prefrontal cortex (PFC) and left inferior frontal gyrus (IFG; P < .001), two in bilateral superior parietal lobe (left: *p* = .002; right: *p* = .040), and one in the superior and middle temporal gyrus around the temporo-parietal junction (*p* < .001).

**Figure 5.**
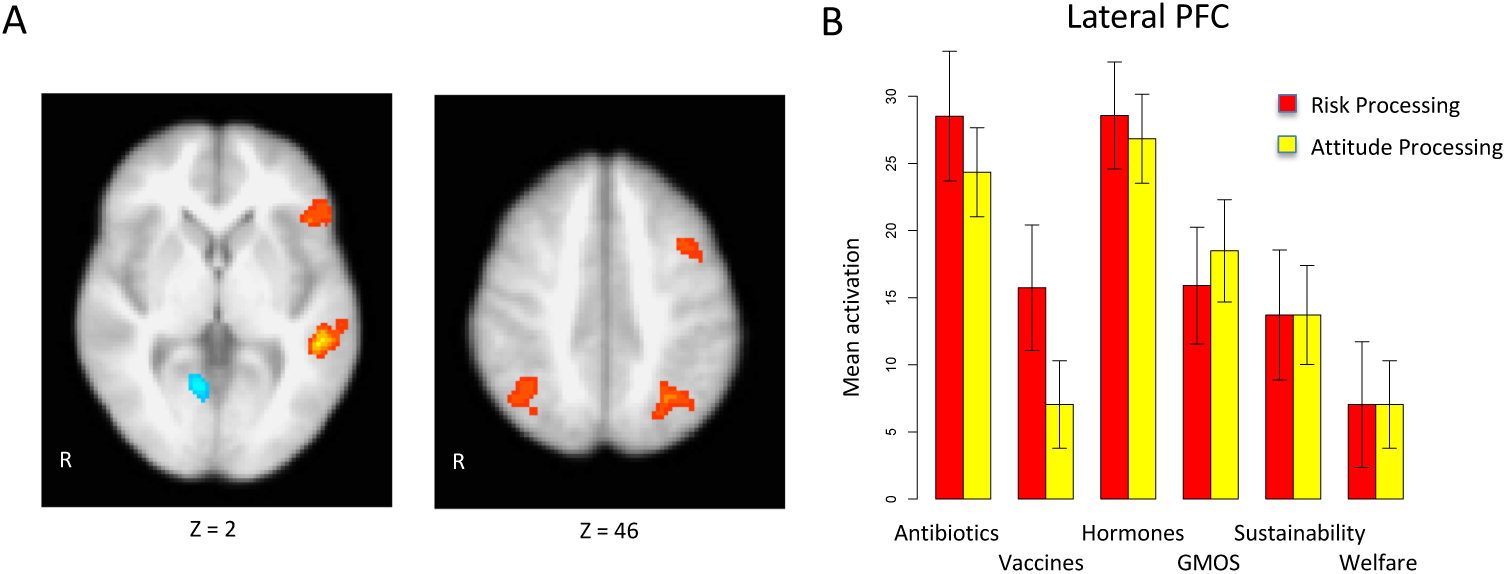
(A) Brain regions positively correlated with between infographic differences in risk perception during the risk processing blocks (red; greater activation = higher risk) and brain regions negatively correlated with between infographic differences in risk during the risk processing blocks (blue; greater activation = lower risk). (B) A bar graph depicting the pattern of activation within the observed lateral PFC cluster. The red bars indicate activation in this region during the risk processing blocks (when participants were asked to process the infographics with respect to risk). The yellow bars indicate activation during the independent attitudes processing blocks (when participants were asked to process attitudes or benefits).

**Table 7.**
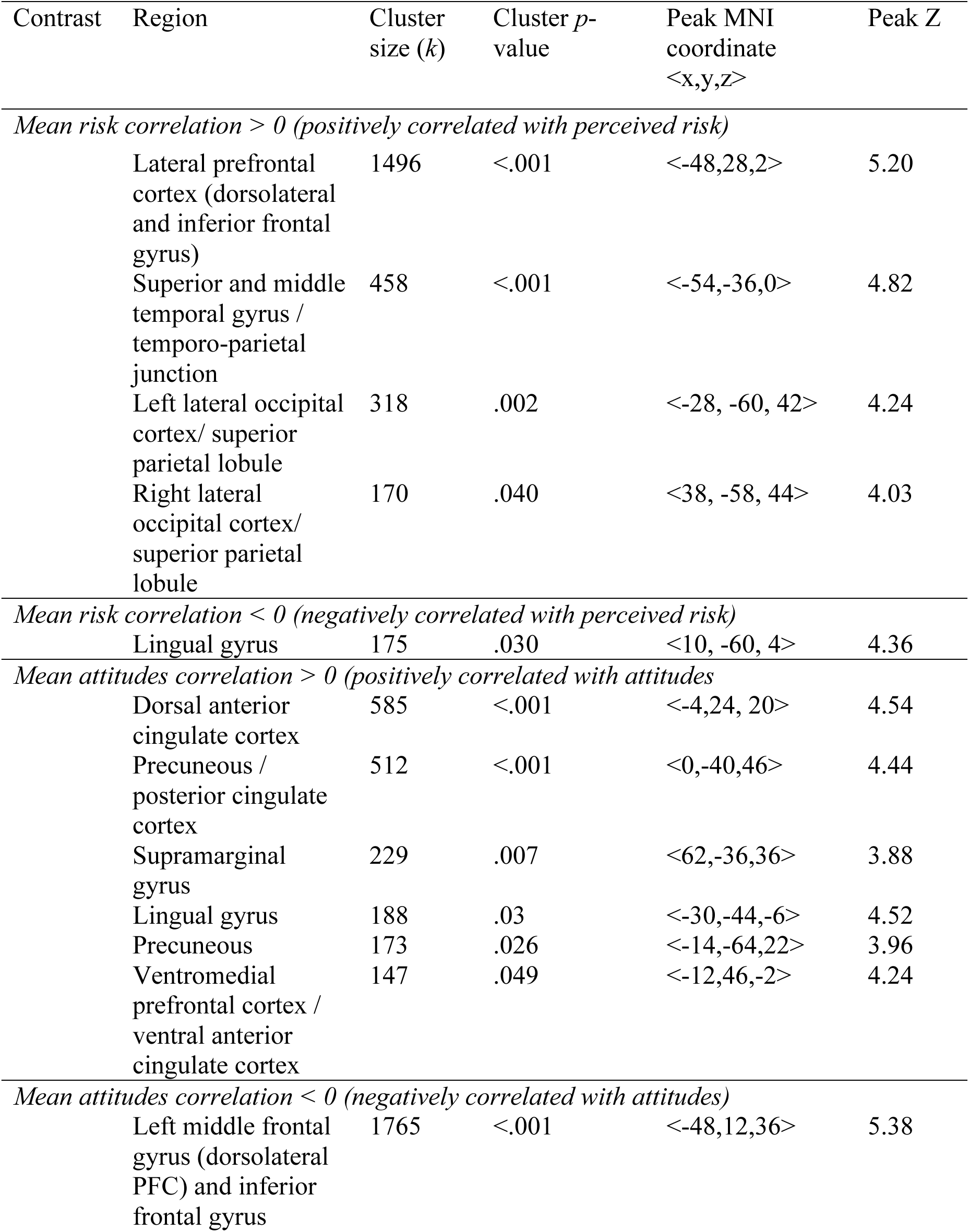
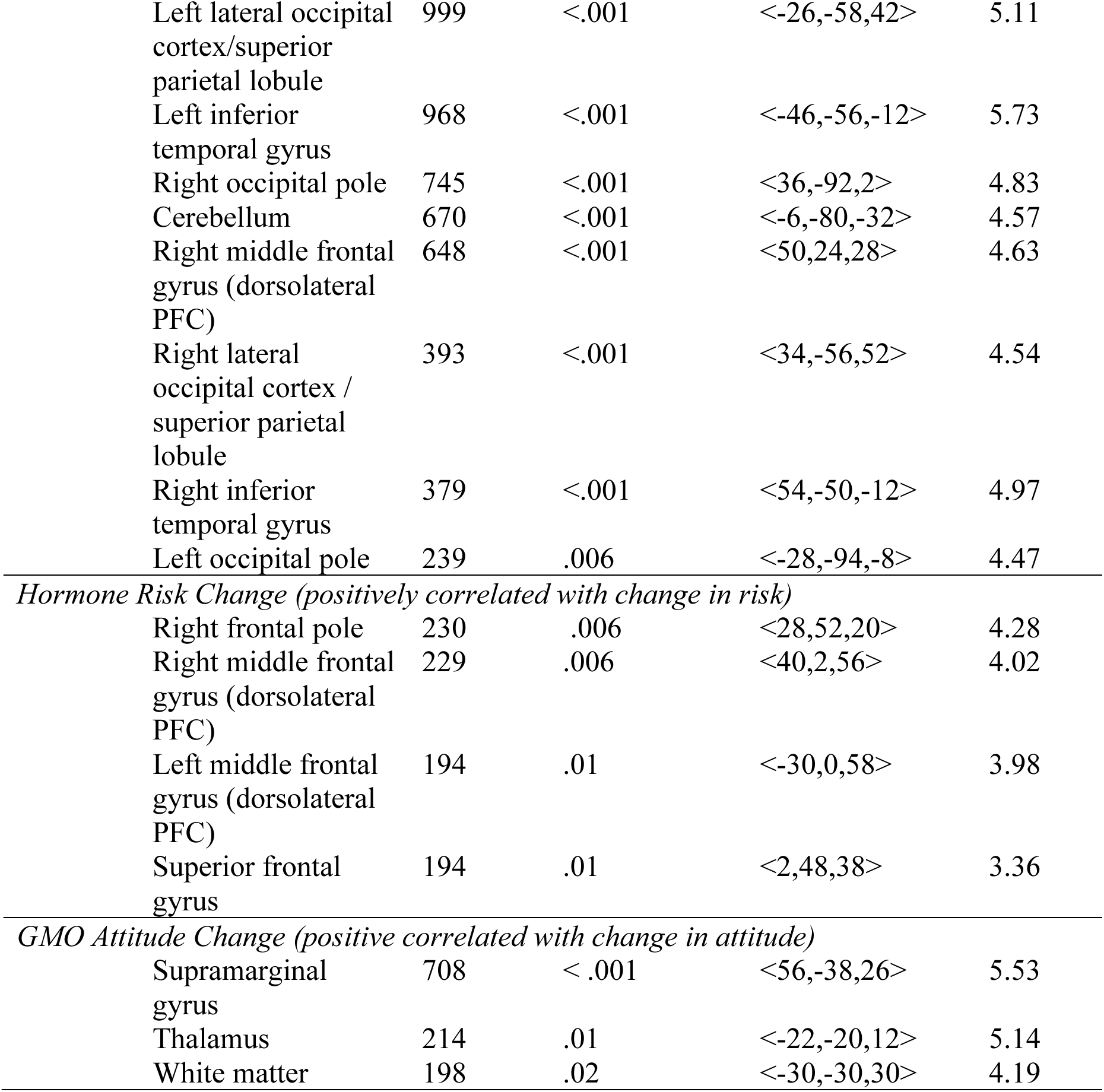
fMRI Results

| Contrast | Region | Cluster size (*k*) | Cluster *p*-value | Peak MNI coordinate<br><*x,y,z*> | Peak *Z* |
| --- | --- | --- | --- | --- | --- |
| *Mean risk correlation > 0 (positively correlated with perceived risk)* |  |  |  |  |  |
|  | Lateral prefrontal cortex (dorsolateral and inferior frontal gyrus) | 1496 | <.001 | <-48,28,2> | 5.20 |
|  | Superior and middle temporal gyrus / temporo-parietal junction | 458 | <.001 | <-54,-36,0> | 4.82 |
|  | Left lateral occipital cortex/ superior parietal lobule | 318 | .002 | <-28, -60, 42> | 4.24 |
|  | Right lateral occipital cortex/ superior parietal lobule | 170 | .040 | <38, -58, 44> | 4.03 |
| *Mean risk correlation < 0 (negatively correlated with perceived risk)* |  |  |  |  |  |
|  | Lingual gyrus | 175 | .030 | <10, -60, 4> | 4.36 |
| *Mean attitudes correlation > 0 (positively correlated with attitudes)* |  |  |  |  |  |
|  | Dorsal anterior cingulate cortex | 585 | <.001 | <-4,24, 20> | 4.54 |
|  | Precuneous / posterior cingulate cortex | 512 | <.001 | <0,-40,46> | 4.44 |
|  | Supramarginal gyrus | 229 | .007 | <62,-36,36> | 3.88 |
|  | Lingual gyrus | 188 | .03 | <-30,-44,-6> | 4.52 |
|  | Precuneous | 173 | .026 | <-14,-64,22> | 3.96 |
|  | Ventromedial prefrontal cortex / ventral anterior cingulate cortex | 147 | .049 | <-12,46,-2> | 4.24 |
| *Mean attitudes correlation < 0 (negatively correlated with attitudes)* |  |  |  |  |  |
|  | Left middle frontal gyrus (dorsolateral PFC) and inferior frontal gyrus | 1765 | <.001 | <-48,12,36> | 5.38 |

|  |  |  |  |  |
| --- | --- | --- | --- | --- |
| Left lateral occipital cortex/superior parietal lobule | 999 | <.001 | <-26,-58,42> | 5.11 |
| Left inferior temporal gyrus | 968 | <.001 | <-46,-56,-12> | 5.73 |
| Right occipital pole | 745 | <.001 | <36,-92,2> | 4.83 |
| Cerebellum | 670 | <.001 | <-6,-80,-32> | 4.57 |
| Right middle frontal gyrus (dorsolateral PFC) | 648 | <.001 | <50,24,28> | 4.63 |
| Right lateral occipital cortex / superior parietal lobule | 393 | <.001 | <34,-56,52> | 4.54 |
| Right inferior temporal gyrus | 379 | <.001 | <54,-50,-12> | 4.97 |
| Left occipital pole | 239 | .006 | <-28,-94,-8> | 4.47 |
| *Hormone Risk Change (positively correlated with change in risk)* |  |  |  |  |
| Right frontal pole | 230 | .006 | <28,52,20> | 4.28 |
| Right middle frontal gyrus (dorsolateral PFC) | 229 | .006 | <40,2,56> | 4.02 |
| Left middle frontal gyrus (dorsolateral PFC) | 194 | .01 | <-30,0,58> | 3.98 |
| Superior frontal gyrus | 194 | .01 | <2,48,38> | 3.36 |
| *GMO Attitude Change (positive correlated with change in attitude)* |  |  |  |  |
| Supramarginal gyrus | 708 | <.001 | <56,-38,26> | 5.53 |
| Thalamus | 214 | .01 | <-22,-20,12> | 5.14 |
| White matter | 198 | .02 | <-30,-30,30> | 4.19 |

The observed lateral PFC cluster is part of the lateral PFC region that we predicted would be associated with between-infographic differences in risk perception. To further illustrate how activation in the lateral PFC varied between the infographics, we extracted mean risk and attitude activation from each subject and plotted them along with the whole-brain results in Figure 5. Note that, because these results are from a region already showing significant tracking of between-infographic risk, it is not appropriate to run additional statistical tests on them (Kriegeskorte, Simmons, Bellgowan, & Baker, 2009); however, the graph is useful to visualize how the lateral PFC is tracking risk. The graph shows that lateral PFC activation tends to be highest for the high perceived risk topics (antibiotics and hormones), but there is less differentiation between the lower perceived risk topics (GMOs, vaccines, animal welfare, and sustainability).

#### 3.2.2 Activation related to attitude processing

Six clusters of activation were positively associated with attitude differences between the infographics (higher activation for more positive attitudes; Figure 6; Table 7), and nine clusters were negatively associated with attitude differences between the infographics (higher activation for more negative attitudes; Figure 6; Table 7). The positively associated regions included the *a priori* region of interest ventromedial prefrontal cortex (*p* = .049), along with dorsal anterior cingulate cortex (ACC; *p* < .001), two clusters in the precuneus (cluster 1: *p* < .001; cluster 2: *p* = .026), the supramarginal gyrus (*p* = .007), and the lingual gyrus (*p* = .030).

The observed ventromedial PFC cluster is part of the ventromedial PFC region that we predicted would be associated with between-infographic differences in attitude positivity. The ventromedial PFC activation is depicted, along with the whole brain activation, in Figure 6. As the bar graph illustrates, this region again primarily distinguished between the less positive attitude (higher perceived risk; antibiotics and hormones) technologies and the more positive attitude technologies. Note that the overall direction of activation is deflected negative in medial PFC, which simply reflects its lower activation relative to the baseline fixation periods. This negative deflection is typical in cognitive tasks as this brain region is part of the “default mode network,” a network of regions that tend to be more active during rest (Raichle, 2015).

**Figure 6.**
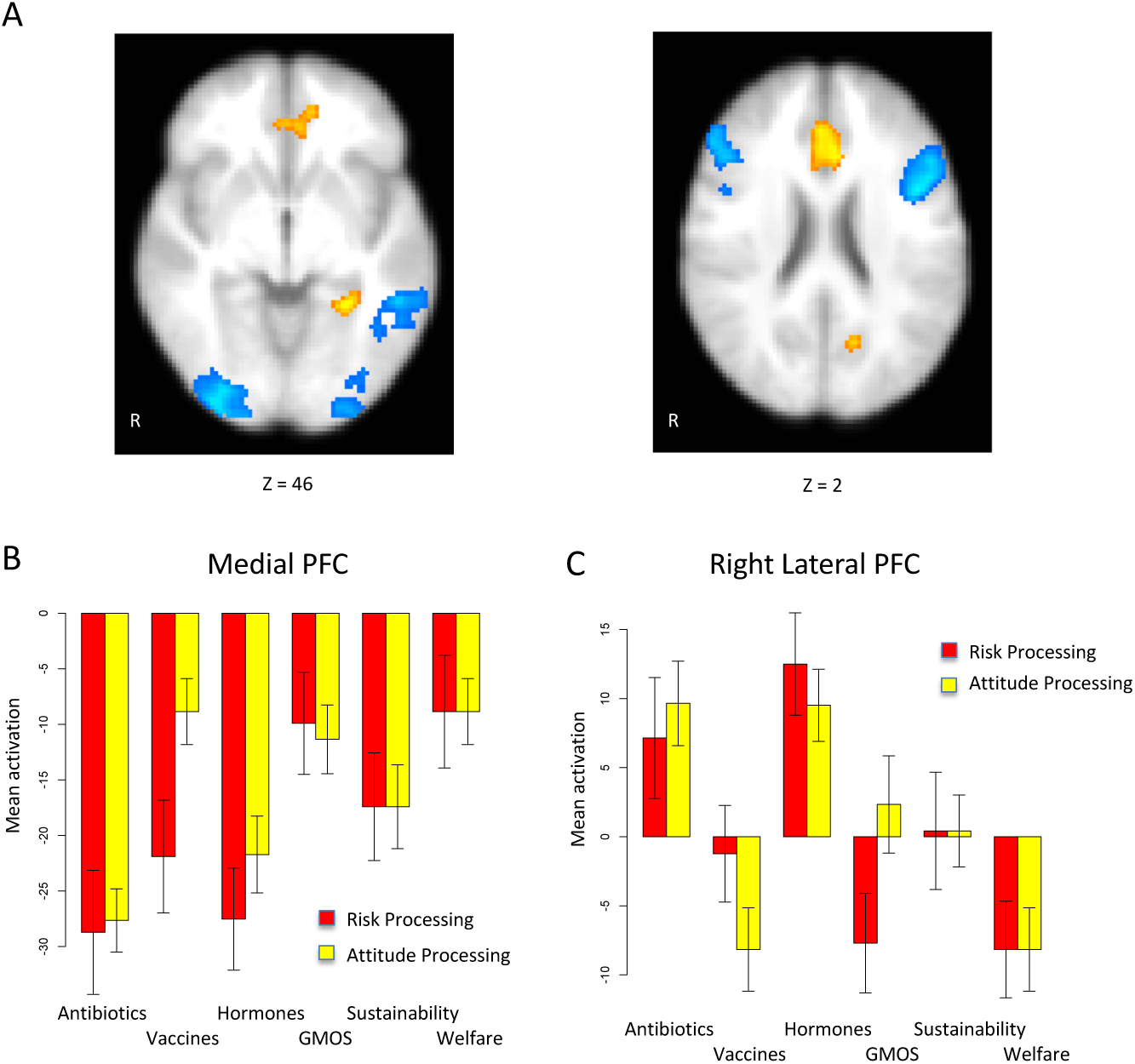
(A) Brain regions positively correlated with between infographic differences in attitudes during the attitude rating blocks (red; greater activation = higher attitudes) and brain regions negatively correlated with between infographic differences in attitudes during the attitude rating blocks (blue; greater activation = lower attitudes). **(B, C)** A bar graph depicting the pattern of activation within the observed medial **(B)** and lateral **(C)** PFC clusters. The red bars indicate activation in this region during the risk processing blocks (when participants were asked to process the infographics with respect to risk). The yellow bars indicate activation during the independent attitudes processing blocks (when participants were asked to process attitudes or benefits).

Clusters that were negatively associated with between-infographic differences in positive attitudes were bilateral (both sides of the brain) lateral PFC regions (both *p* < .001), the left of which overlaps with the cluster revealed for the risk perception analysis, as well as superior parietal regions that overlapped with the risk results (both *p* < .001). The lateral PFC region was again part of predicted lateral PFC region that we expected to be negatively associated with attitudes (more activation for less positive attitudes).

#### 3.2.3 Individual Differences in Attitude Change

In addition to examining overall risk and attitudes associated with the infographics, we wanted to examine whether any brain regions were predictive of an individual’s change in attitudes or risk perception toward the food technologies. To this end, we entered participants’ post - pre-infographic attitude and risk perception Z change scores into third-level whole-brain GLM analysis to test which brain regions were significantly associated with individual differences in risk and attitude change.

Two of these analyses showed significant results: change in risk perception for the hormone infographic and change in attitudes for the GMOs infographic. For the hormone infographic (Figure 7A & 7C; Table 7), we found four clusters that were significantly associated with change in risk perception: bilateral lateral PFC (Left: *p* = .01; Right: *p* = .006), superior frontal gyrus (*p* = .01), and frontal pole (*p* = .006). Specifically, these regions were positively associated with change in risk perception, which indicates that these regions were associated with either resisting change in risk (as most participants decreased) or increasing perceptions of risk. These results help to interpret the within-subject lateral PFC effects, where the lateral PFC was associated with higher risk perception and less positive attitudes across the infographics. Specifically, they suggest that the lateral PFC may be inhibiting attitude change for hormones, as opposed to encouraging attitude change. However, these results seem to be specific for hormones, as they did not appear for any of the other risk change measures.

**Figure 7.**
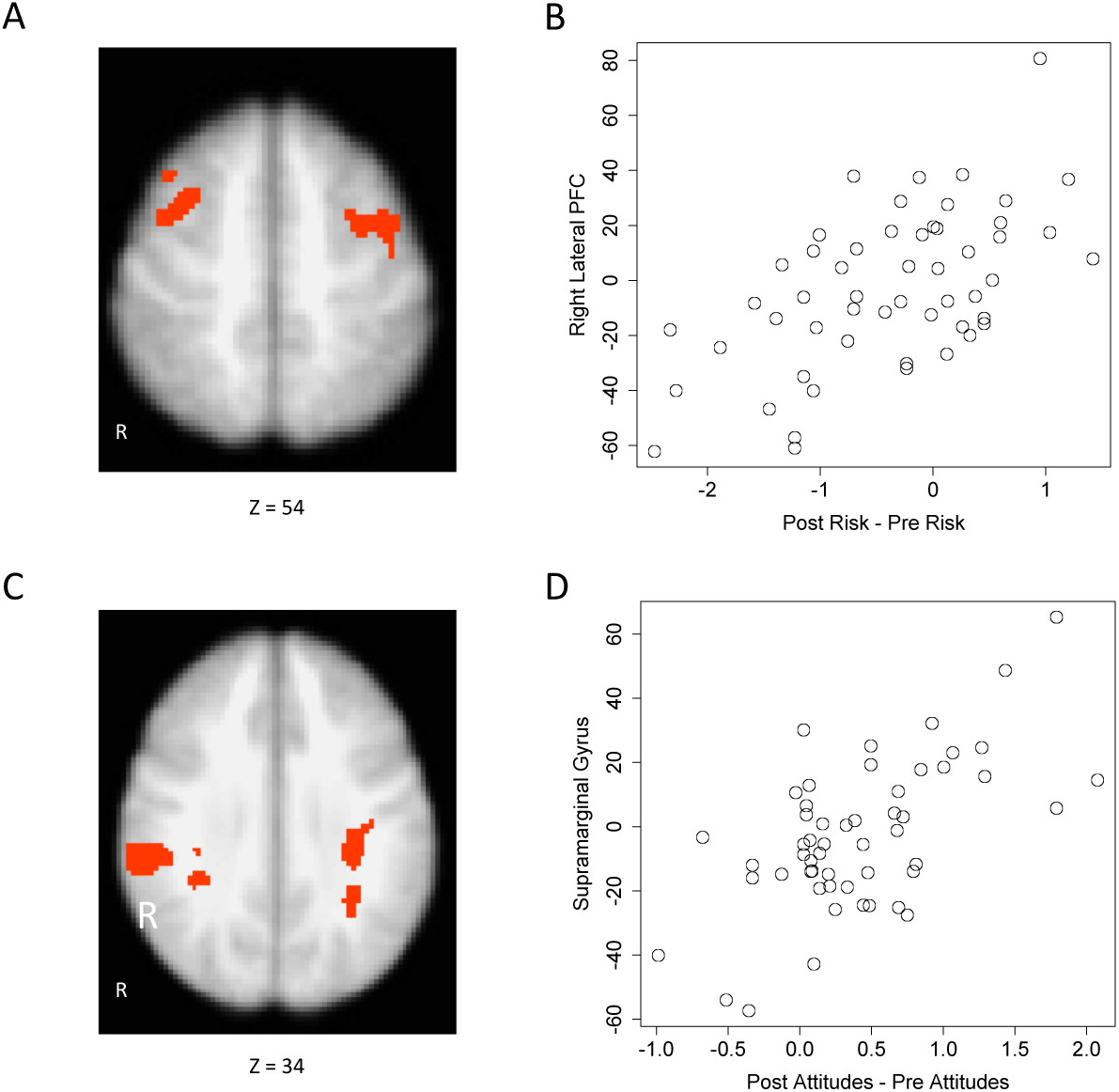
**(A)** Results from the whole brain analysis identifying areas that are positively correlated with change in risk perception for hormones. **(B)** Scatterplot illustrating association between the right lateral PFC and change in risk perception for hormones. **(C)** Results from whole brain analysis identifying areas that are positively associated with change in attitudes for GMOs. **(D)** Scatterplot illustrating association between right supramarginal gyrus and change in attitudes for GMOs. Note that the scatterplots in these figures are primarily suitable for evaluating the relationship (e.g., the direction of association, linearity assumption) depicted in the brain

In addition to the hormone results, we found three significant clusters that were positively associated with attitude change for GMOs (higher activation for more positive attitude change): in the supramarginal gyrus (*p* < .001), the thalamus (*p* = .01), and left cerebral white matter (*p* = .02; Figure 7B & 7C; Table 7).

## 4. Discussion

Food technologies play an important role in food quality, sustainability, and global food security, yet technological innovation is often met with resistance from consumers. The present study used neuroimaging to reveal how the brain processes information related to food technology topics and how these neurobiological processes vary with perceived riskiness and attitudes associated with the technologies. Participants viewed infographics about six different food technology topics including antibiotics, hormones, GMOs, vaccines, sustainability, and animal welfare. We found that activation in lateral PFC tracked between-infographic differences in risk perception and attitudes and was most activated for the higher perceived risk / less positive attitude topics (antibiotics and hormones). In contrast, ventromedial PFC activation tracked positive attitudes and was most activated for the topics (GMOs, vaccines, sustainability, and animal welfare) associated with more positive attitudes. In addition to our primary, between-infographic results, we found individual differences in risk perception change for hormones was negatively correlated with lateral PFC activation, suggesting that, for this topic, higher lateral PFC activation was associated with resistance to change in risk perception. Finally, part of the supramarginal gyrus was positively correlated with individual differences in attitude change for GMOs.

The fMRI results present a critical step forward in understanding consumer attitudes and risk perceptions surrounding food technology. Because education and dissemination of information are necessary tools for reducing consumer resistance to technological development, it is important to understand how processing such information is related to differences in attitudes and risk perception. Functional neuroimaging provides a promising window into consumer information processing as it has the potential to show—above and beyond self-report surveys—which information is being processed overall more positively, by examining how activation in regions associated with positive subjective value, such as the ventromedial PFC, track attitudes and risk perception. It also has the potential to uncover which information is being processed more effortfully, which tends to activate regions associated more with uncertainty processing. Brain activation patterns can thus reveal how and whether informational campaigns are leading to effective processing by consumers, and potentially index attitude and risk perception change.

Despite the success of the current study at identifying regions that track attitude and risk perception differences between food technologies, it is important to note that it remains unclear whether attitudes are affecting how the information about the technologies is being processed or are a reflection of the information the infographics contain. For example, the increased ventromedial PFC may reflect more positive attitudes toward animal welfare technologies and sustainability in general, or may reflect the fact that participants like the specific contents of these infographics better. Future research will need to dissociate individuals’ baseline affective responses to and cognitive information about food technology topics from the information present in the infographics and determine how these factors overlap. Nevertheless, the current results point to connections between infographic processing and attitudes/risk perception that are useful for understanding how consumers approach information about food technologies.

The present results complement and extend previous fMRI studies on willingness-to-pay for foods (e.g., milk, eggs) produced with food technologies. Previous studies have largely focused on the same lateral PFC regions that we identified here. The lateral PFC tends to be more active when participants are considering both price and technology in willingness-to-pay judgments, in addition to tracking participants’ choices to opt for more expensive organic or natural choices (Linder et al., 2010; Lusk et al., 2015). These results are related to the present finding that the lateral PFC tends to activate more for the less liked, higher perceived risk technologies and may reflect the same general uncertainty-related mechanism. When people are confronted with technologies that conflict with their preferences, the lateral PFC may be engaged to help them resolve this uncertainty. Likewise, consistent with previous findings that the ventromedial PFC tracks the outcomes of willingness-to-pay judgments (Crespi et al., 2015), we found the ventromedial PFC tracks overall attitudes for the technologies.

What is critical about the present results, relative to previous willingness-to-pay studies, is that such lateral PFC and medial PFC activation are connected to information processing surrounding technologies outside of choice-related contexts. This suggests that the activation found in previous willingness-to-pay studies may not reflect simple trade-offs between price and technology, but more basic pre-decisional attitudes that feed into such choice trade-offs. For example, it may be the case that lateral PFC activation during a choice to pay more for organic foods specifically reflects less positive attitudes toward technologies as opposed to uncertainty about the decisions per se. Whether the PFC activation observed in previous studies reflects an attitudinal “input” into the decision process or an “outcome” of the decisional process (see e.g., Davis, Love, & Maddox, 2009) is an important question for future research. In many cases, attitudes, risk perception and decision processes are tightly coupled, and thus the practical upshot—that people will pay to avoid technologies that generate conflict or uncertainty—may be the same in both scenarios.

In addition to its higher activation for technologies that were perceived as higher risk and associated with less positive attitudes, our finding that lateral PFC activation was negatively correlated with individual differences in change in risk perception for hormones gives important information about the lateral PFC’s potential function in the present context. The lateral PFC has been shown in previous studies to track individual differences in decision-making and cognitive performance (Christopoulos et al., 2009; Schonberg et al., 2011; Worthy et al., 2016). Although the lateral PFC has been shown to track participants’ willingness to pay more to avoid food technologies and is often associated with individual differences in risk aversion, other results suggest that activation in lateral PFC may not always be associated with increases in negative attitudes or perceived risk. Indeed, a recent study on willingness to pay for animal welfare technologies in egg production has shown that lateral PFC may more generally reflect updating of willingness-to-pay based on information (McFadden, et al., 2015). This could mean that lateral PFC activation is not necessarily positively or negatively related to acceptance of a technology, but rather simply reflects controlled deliberation (i.e., elaboration; Petty & Cacioppo, 1986). The outcome of such controlled deliberation may be an increase or decrease in attitudes (or potentially even no change), depending on several factors such as argument strength, personal relevance, cognitive ability, and participants’ motivations for changing their attitudes (Petty, Briñol, & Priester, 2009). For example, here lateral PFC was associated with resistance to changing risk perception for hormones, but perhaps with different arguments or with participants motivated to change their risk perceptions, it may have been associated with decreased risk perception.

Interestingly, in the present study, the results for risk perception and positive attitudes were largely overlapping in the sense that the same lateral PFC regions that tracked between-infographic differences in risk perception also tended to be negatively correlated with positive attitudes (higher activation for less positive attitudes). These results suggest, in the present case, that risk perception and attitudes may tap many of the same neurobiological systems. This is not a surprise as risk perception is often tightly coupled with attitudes, and attitudes are a facet of risk perception (Kraus, Malmfors, & Slovic, 1992). However, whether this observation will extend to other areas of risk perception or attitudes remains an open question. One reason why the present results may overestimate the neurobiological overlap between risk perception and more general attitude processing is that many of the technologies were associated with low risk overall. Indeed, only the highest perceived risk categories (antibiotics and hormones) consistently exceeded the scale midpoint (2.5) and even then, they were well below the limits of the scale (scale max = 5; see Table 2). It is possible that more extreme risks may tap additional brain regions associated with fear and negative affect, such as the amygdala (Feinstein, Adolphs, Damasio, & Tranel, 2011).

A second reason we may not have observed large differences between the brain regions tracking attitudes and risk perceptions is that the infographics chosen for the study were developed as tools for promoting food technologies and thus did not contain extensive risk information (e.g., of possible antibiotic resistance from livestock use). Although some of the infographics presented information relevant to aspects of chemical risk perception, such as dose-to-response reasoning (e.g., in the hormones and antibiotics infographics; Kraus, Malmfors, & Slovic, 1992), or risks to livestock from wild animals or the elements (animal welfare infographics), none of the information would be expected to increase risk perception *a priori*. Our goal in the present study was not to induce perception of risk *per se* by presenting risky information, but instead to test how brain activation tracked people’s general perceptions of risk about the technologies, and how such risk perception related to infographic processing. In this way, our study was consistent with the majority of risk perception studies on chemicals and food technologies that simply seek to measure people’s perceptions of risk without telling them specific known risks of such chemicals and technologies (e.g., Cox & Evans, 2008; Kraus, Malmfors, & Slovic, 1992). Future studies should consider testing whether presenting participants with information intended to increase perceptions of risk would change which brain regions track risk perception. Indeed, a mix of information that is congenial and uncongenial to participants’ baseline inclinations may better represent the real-life information environment and provide opportunities to study the roles of selective information search and memory on perceptions of food technologies (Hart et al., 2009).

With respect to whether brain regions involved in risk perception and attitude processing may diverge in studies using riskier technologies or those giving participants more risk-specific information, another possible limitation of the current study is that participants may not have followed the cue to focus on risks or attitudes/benefits during infographic processing at all. For example, it is possible participants focused on the same information in the infographics regardless of whether they were cued to focus on risks or personal attitudes/benefits. As we discuss above, the cueing procedure was primarily intended to encourage active processing of the infographics and prime participants for what types of questions they would be answering next. Because all participants received the same cues, it is not possible to definitively answer whether this priming changed how participants processed the infographics. Indeed, the strong consistency of the results across cue types suggests that the participants may not have changed their infographic processing substantially based on the cues. However, the individual difference results are suggestive that some risk and attitude-specific processing differences across cue types may have occurred. For example, lateral PFC activation for hormones during the risk perception blocks specifically correlated with individual differences in risk perception change for hormones, but this association was not observed for attitude change during the attitude change blocks. Nevertheless, our primary goal of examining how risk perception and attitudes relate to brain processing during infographic viewing is not dependent on participants changing how they processed the infographics based on the cues. For example, even if all infographic viewing blocks were processed the same (or if there were no cues presented at all), our findings that brain activation during infographic processing tracks subsequent between-infographic differences in risk perception and attitudes presents a useful starting point for studying how information about food technologies is processed by the brain.

A final limitation of the present study is that the infographics we employed contained a wide range of varied strategies to convey the safety and utility of the food technologies. Thus, it is not possible to separately conclude which aspects of the infographics may have been stronger or weaker in influencing attitudes and risk perception, or brain activation, the most. For example, the hormones infographic contains information highlighting the low tissue accumulation of hormones from hormone implants, relative to other dietary sources of estrogen, information about sustainability and environmental benefits, and the fact that the implants replace some naturally occurring hormones that are removed by food production practices. Any of these strategies could be leading to the observed attitude and risk perception results and activation for this infographic. Future studies interested specifically in isolating cognitive and neurobiological factors that lead to increases in attitudes and consumer acceptability may wish to constrain the arguments to be presented in more serial fashion to isolate their individual effects. Nevertheless, the use of preexisting real-world infographics offers a strong, ecologically valid simulation of how participants may encounter information about food technologies outside of the laboratory. The tight coupling we observed between participants’ attitudes and risk perception and brain activation suggests that fMRI may be useful for screening such infographics before public information campaigns are deployed.

## 4. Conclusions

The present study examined the neurobiological processes engaged when participants viewed information related to food technologies. We found that areas of the lateral PFC tracked heightened risk perception and less positive attitudes for technologies, whereas the medial PFC tracked positive attitudes. These results give a unique window into the minds of consumers, as they illustrate how differences in attitudes and risk between food technologies affect basic information processing in the brain. Connections between the basic attitudinal-risk processing and individual differences in activation suggest that functional neuroimaging may be a useful tool for assessing how people process informational materials about the benefits of food technologies, which may lead to more effective strategies for combatting resistance to such technologies. Indeed, to promote increasing advances in food quality, sustainability, and security, technological innovation will continue to be critical, and consumer acceptance of such innovation will continue to play a large role in determining whether it succeeds.

## ACKNOWLEDGEMENTS

This study was funded by Merck Animal Health grant PO7100310693 to T.D. and M.F.M. M.F.M. has a consulting relationship with and J.L.F. is employed by Merck Animal Health. M.F.M. and J.L.F. were involved in the study design but did not collect, analyze, or interpret the data. All authors were involved in writing of the report, and the decision to publish the manuscript. Address correspondence to.

## Supplementary Material

**Supplemental Table 1.**
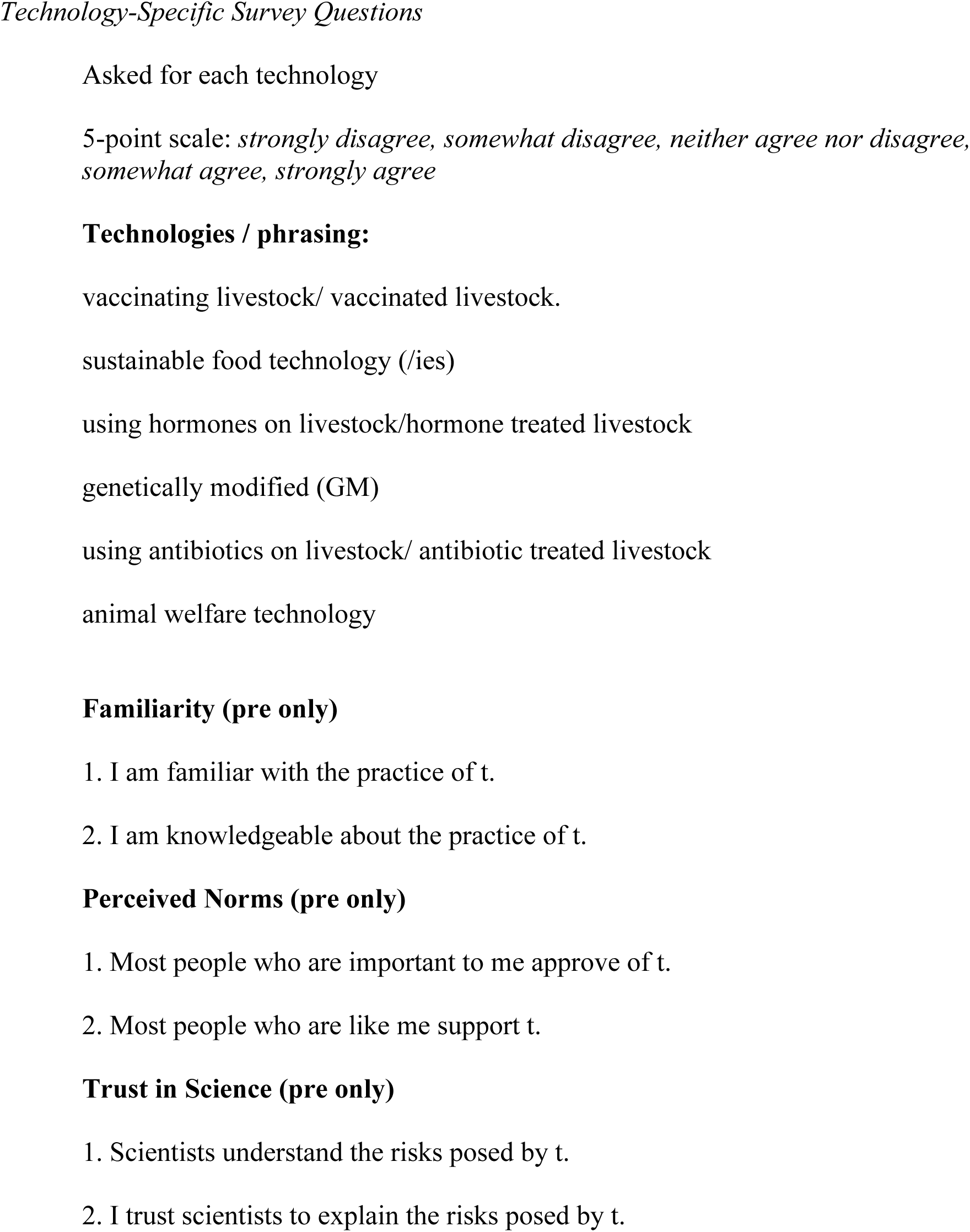
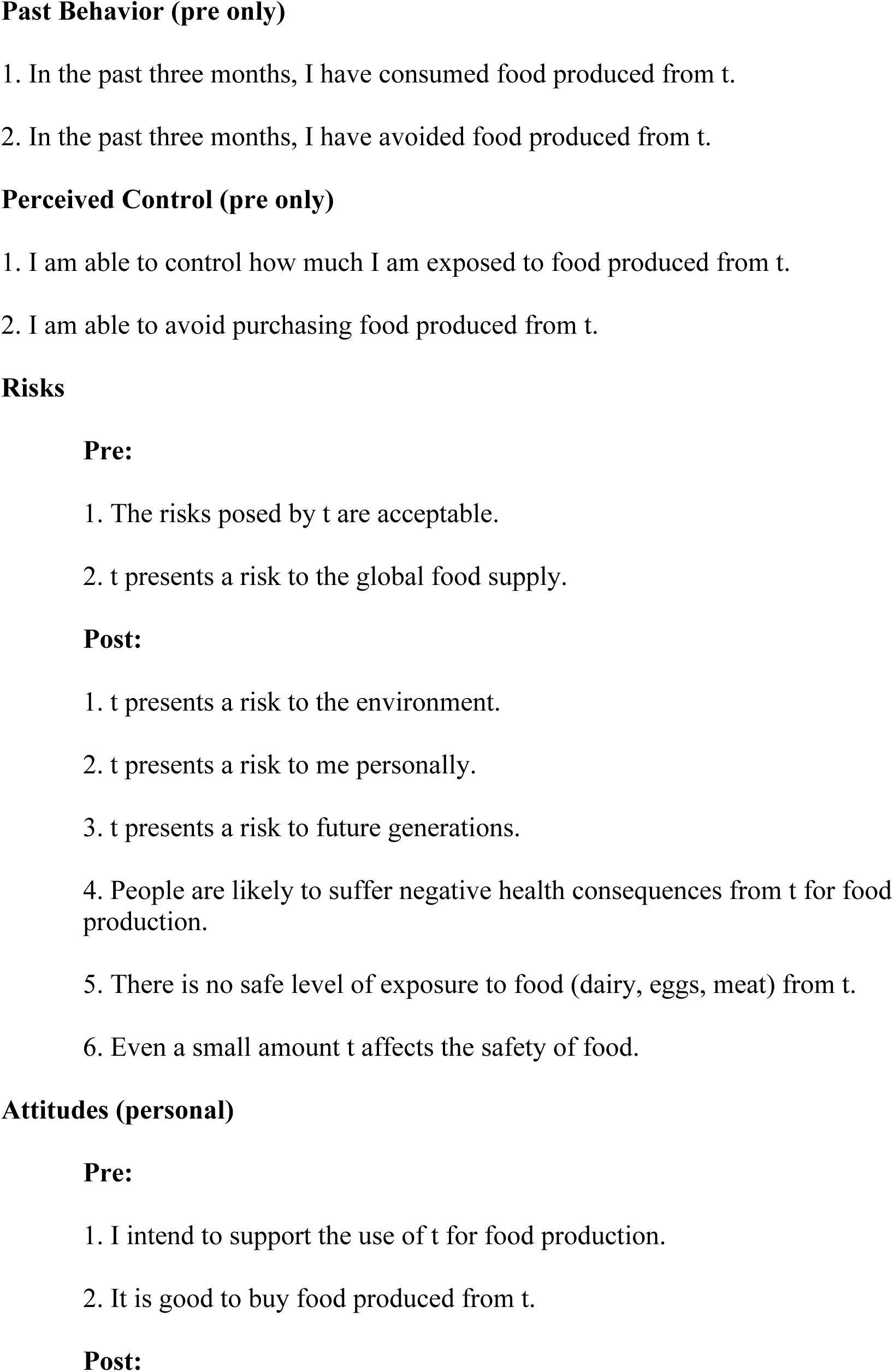
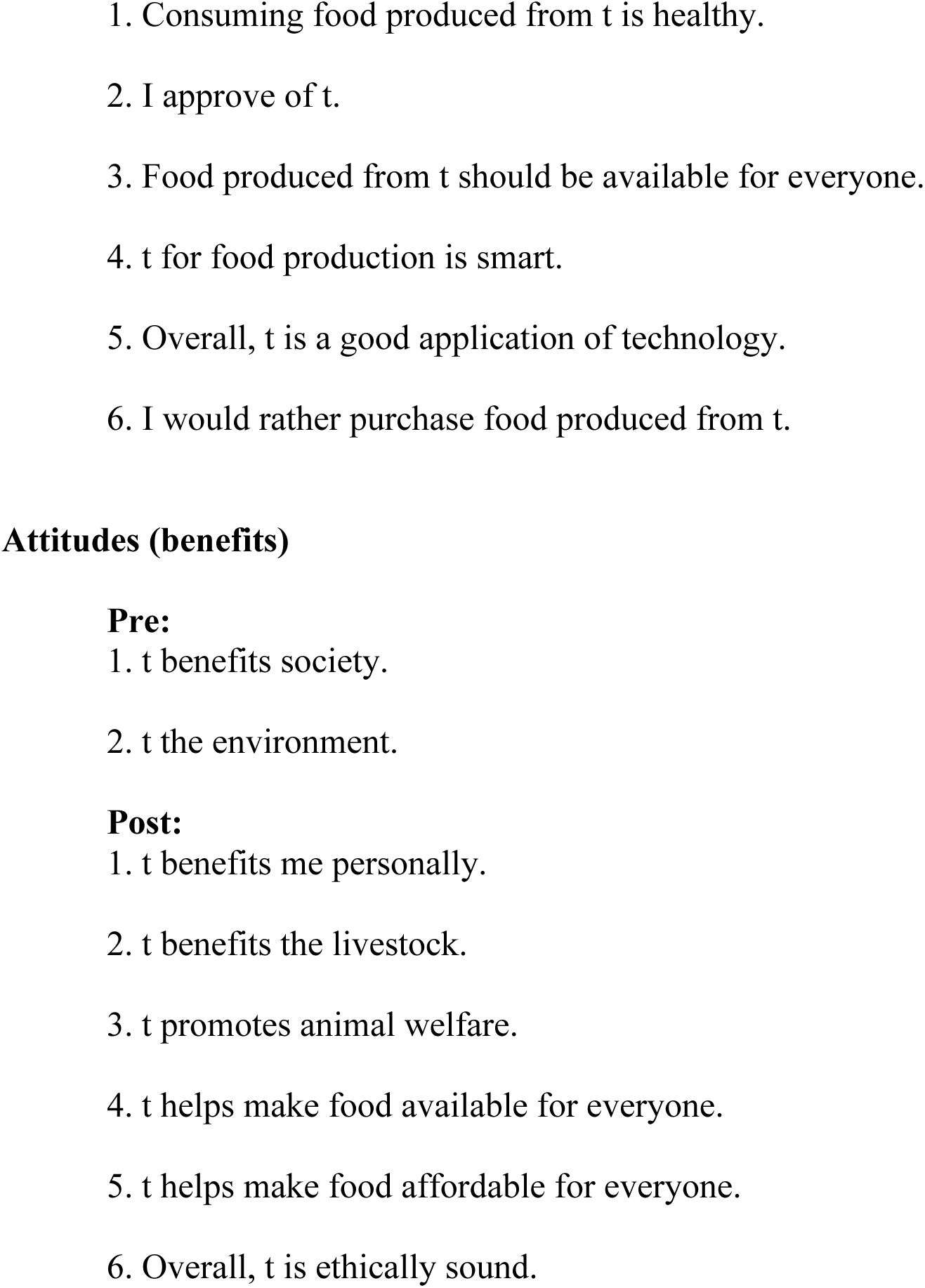
Wording of the attitudes and risks questions were based on typical risk perception (Slovic, 2016) and Theory of Planned Behavior survey construction (Ajzen, 1991).

**Supplemental Table 2.** Cronbach’s coefficient alpha for each survey measure from the online norming study.

| Measures | $\alpha$ |
| --- | --- |
| *Antibiotics* |  |
| Risk (pre) | .82 |
| Risk (post) | .93 |
| Attitudes (pre) | .94 |
| Attitudes (post) | .96 |
| Familiarity | .84 |
| Norms | .90 |
| Trust in science | .81 |
| Behavior | .62 |
| Control | .93 |
| *Vaccines* |  |
| Risk (pre) | .83 |
| Risk (post) | .94 |
| Attitudes (pre) | .91 |
| Attitudes (post) | .96 |
| Familiarity | .89 |
| Norms | .89 |
| Trust in science | .85 |
| Behavior | .55 |
| Control | .89 |
| *Hormones* |  |
| Risk (pre) | .75 |
| Risk (post) | .94 |
| Attitudes (pre) | .93 |
| Attitudes (post) | .96 |
| Familiarity | .85 |
| Norms | .90 |
| Trust in science | .80 |
| Behavior | .72 |
| Control | .89 |
| *GMO* |  |
| Risk (pre) | .88 |
| Risk (post) | .96 |
| Attitudes (pre) | .94 |
| Attitudes (post) | .97 |
| Familiarity | .82 |
| Norms | .89 |
| Trust in science | .85 |
| Behavior | .67 |
| Control | .88 |
| *Sustainability* |  |
| Risk (pre) | .78 |
| Risk (post) | .92 |
| Attitudes (pre) | .93 |
| Attitudes (post) | .95 |
| Familiarity | .93 |
| Norms | .91 |
| Trust in science | .81 |
| Behavior | .38 |
| Control | .89 |
| *Animal welfare* |  |
| Risk (pre) | .74 |
| Risk (post) | .91 |
| Attitudes (pre) | .87 |
| Attitudes (post) | .94 |
| Familiarity | .94 |
| Norms | .88 |
| Trust in science | .80 |
| Behavior | .35 |
| Control | .87 |

**Supplemental Table 3.**
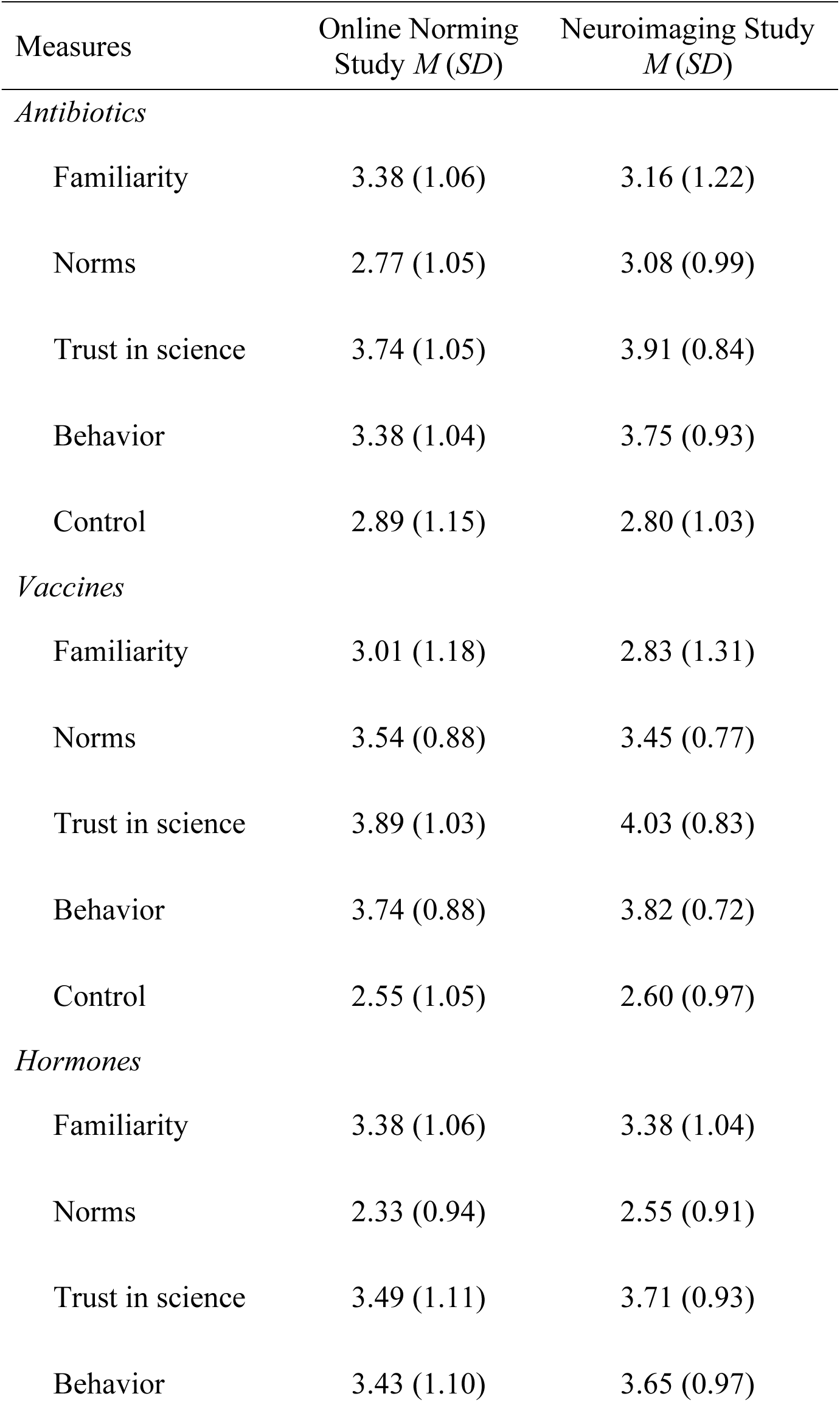
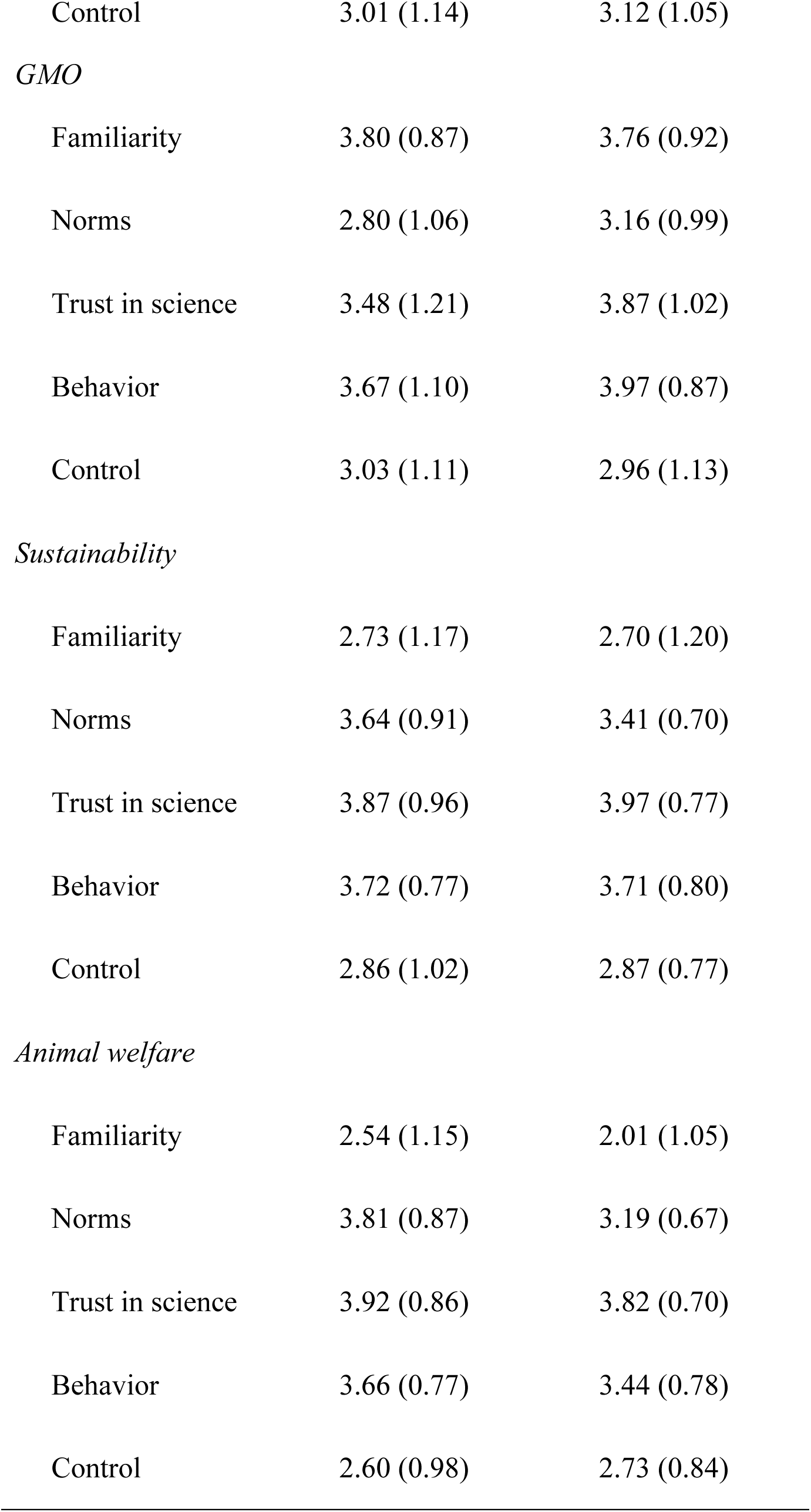
Means and standard deviations of survey measures for the online norming study and the neuroimaging study.

**Supplemental Table 4.** Mean, standard deviation, and reliability statistics (standardized alpha) for standardized survey measures. Note: Standardized survey measures were taken after seeing infographics in the neuroimaging study.

|  | <u>Online</u><br><u>Norming</u><br><u>Study</u> | <u>Neuroimaging</u><br><u>Study</u> |
| --- | --- | --- |
| Measure | *M (SD)* | *M (SD)* |
| Food Tech.<br>Neophobia | 3.04 (0.85) | 2.77 (0.62) |
| Trust in Science | 3.92 (0.87) | 4.04 (0.65) |
| Chemical risk<br>perception: Dose-to-response<br>reasoning | 3.36 (0.77) | 3.25 (0.70) |
| Chemical risk<br>perception:<br>Attitudes toward<br>chemicals | 3.26 (0.86) | 2.96 (0.60) |
| Chemical risk<br>perception:<br>Chemical risk<br>reduction | 2.62 (0.86) | 2.59 (0.63) |
| Chemical risk<br>perception:<br>Miscellaneous | 2.86 (0.52) | 2.89 (0.40) |

**Supplemental Figure 1.**
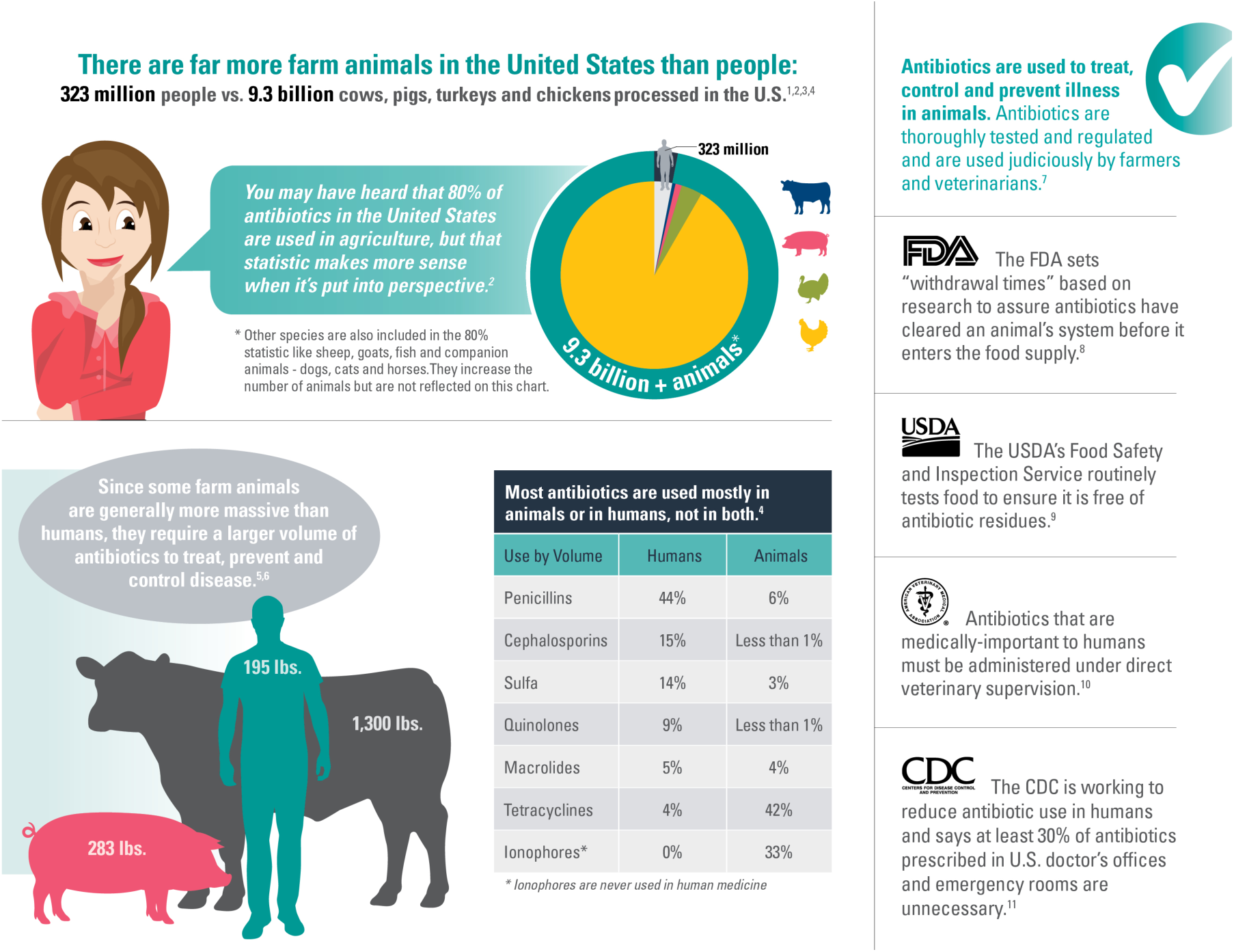
Antibiotics infographic used in the study.

**Supplemental Figure 2.**
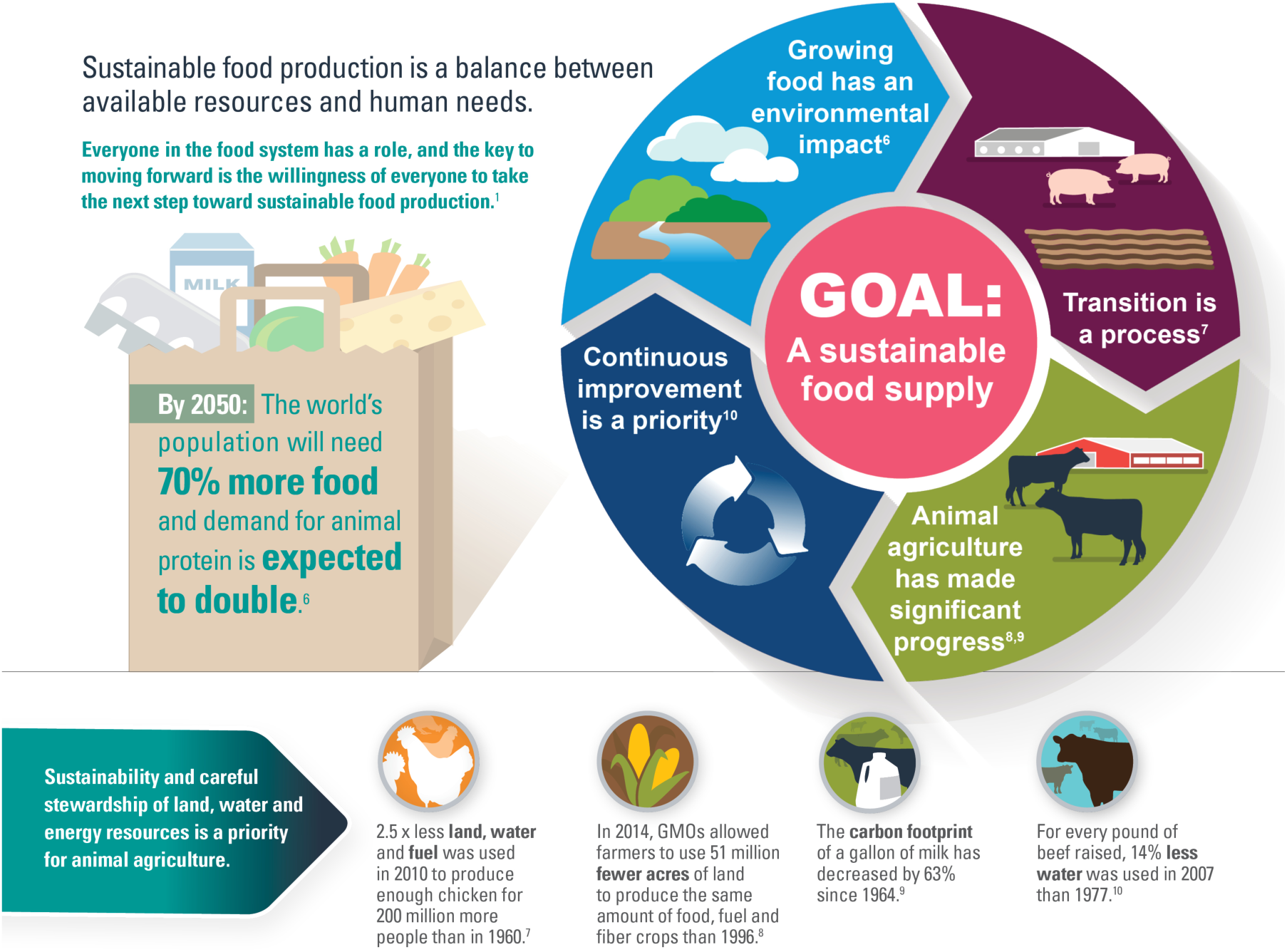
Sustainability infographic used in the study.

**Supplemental Figure 3.**
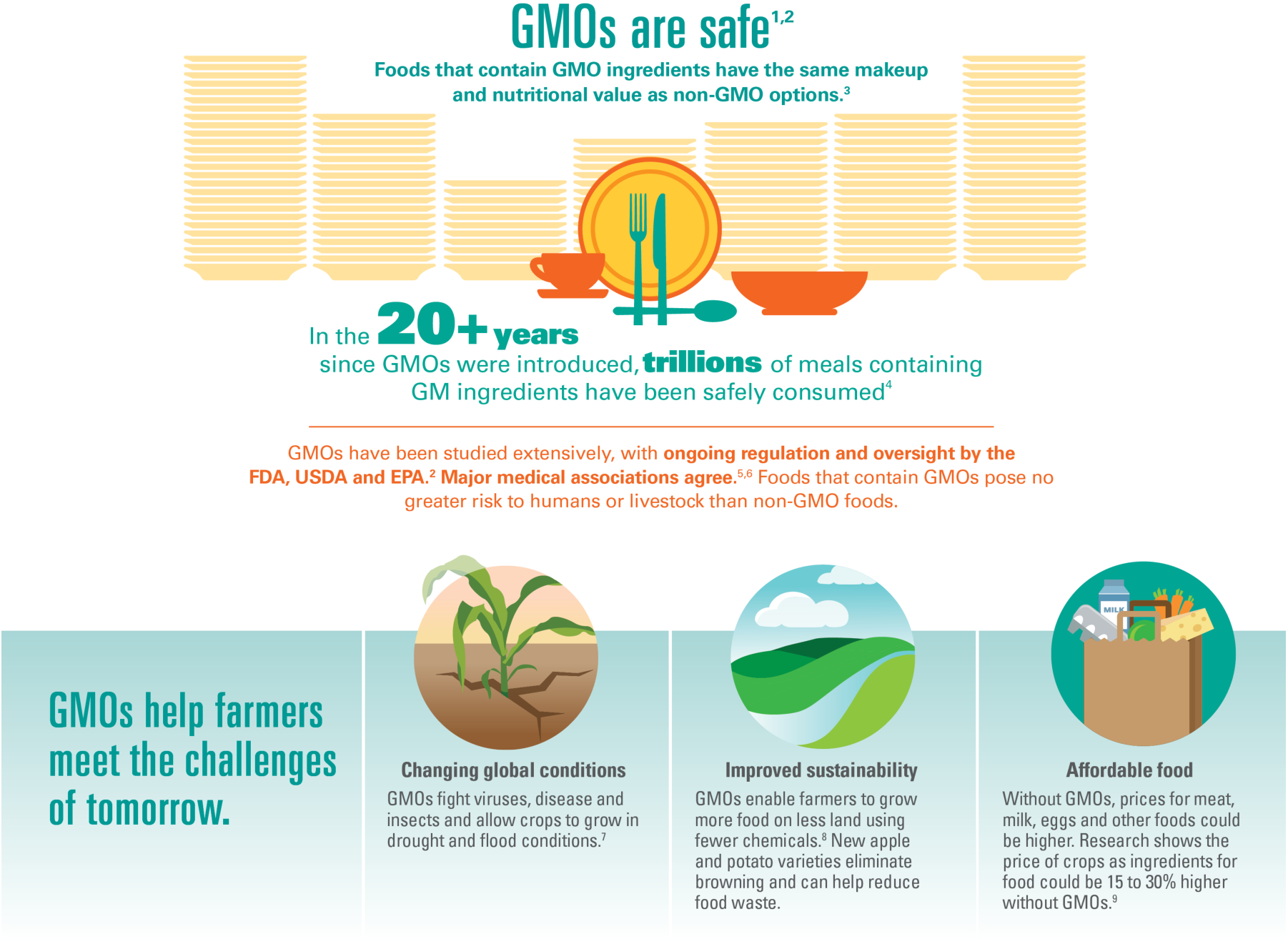
GMO infographic used in the study.

**Supplemental Figure 4.**
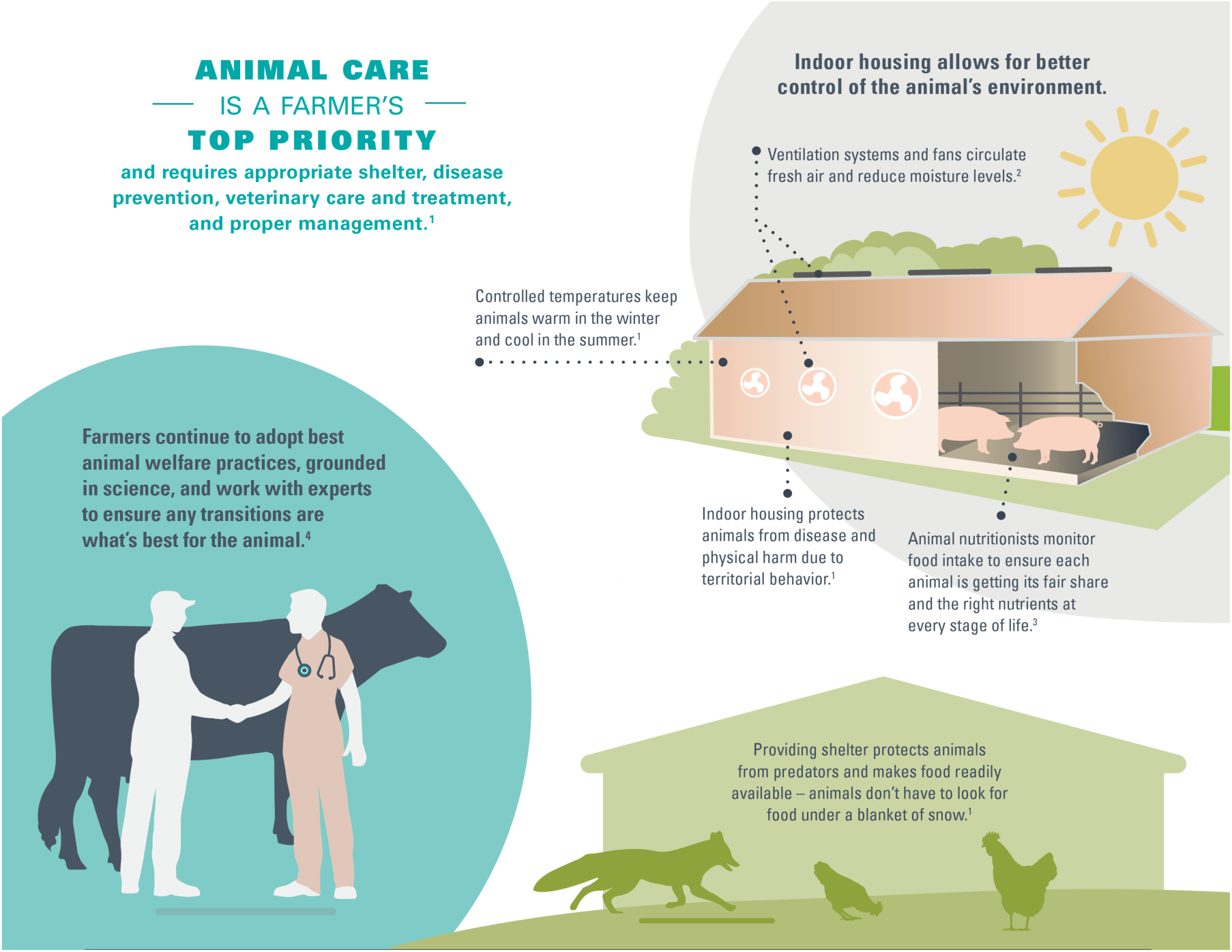
Animal welfare infographic used in the study.

**Supplemental Figure 5.**
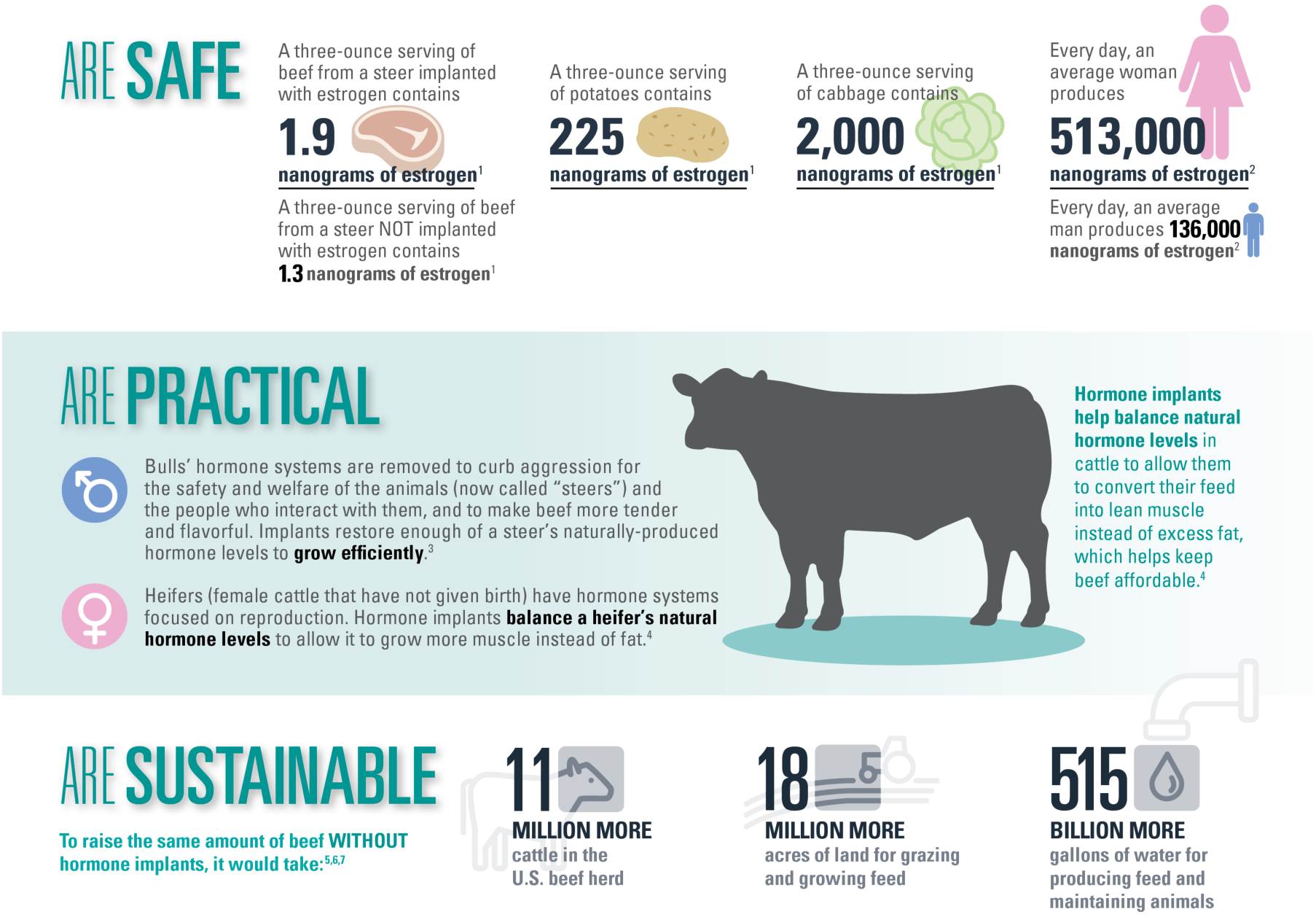
Hormone infographic used in the study.

**Supplemental Figure 6.**
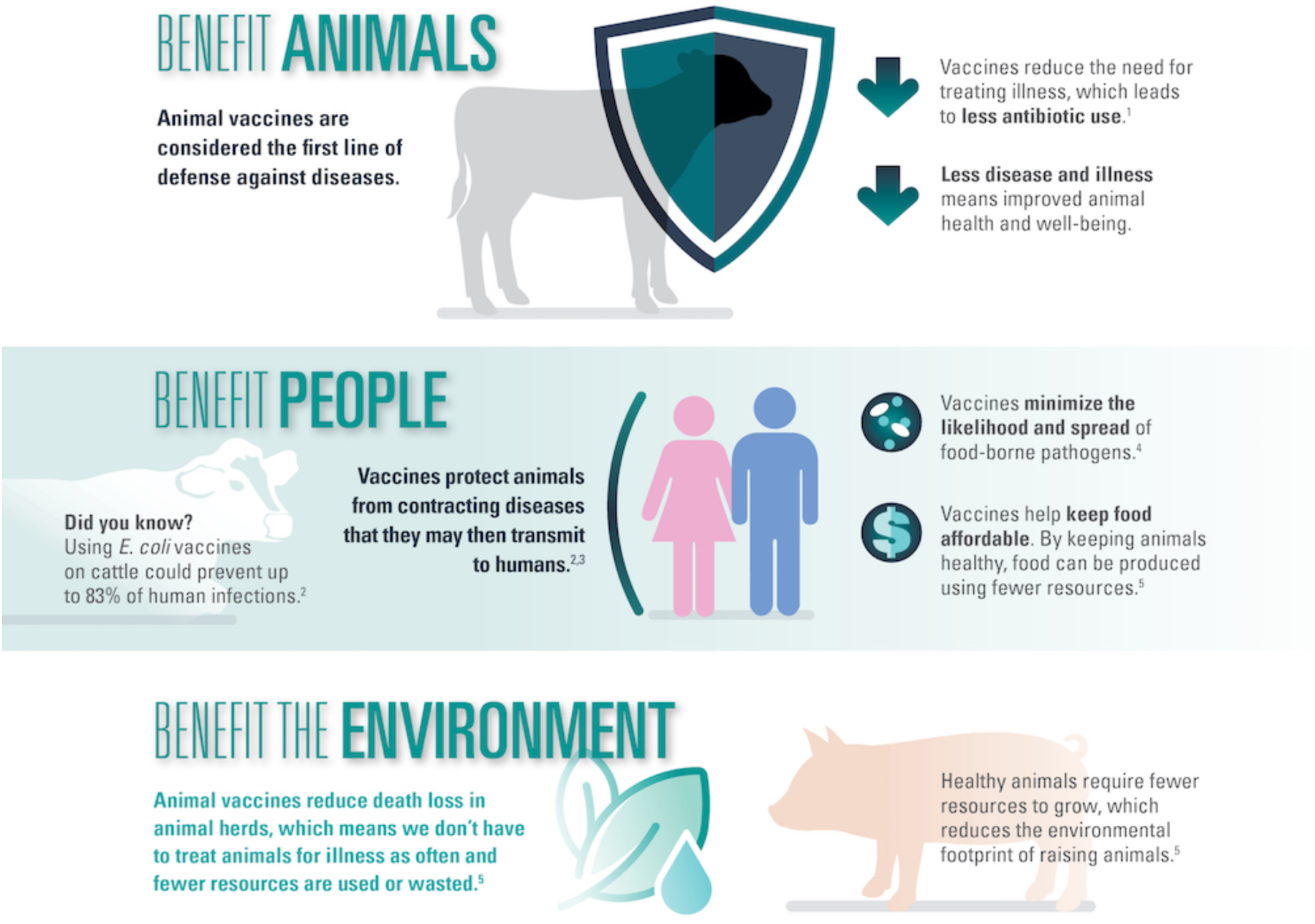
Vaccine infographic used in the study.

**Supplemental Figure 7.**
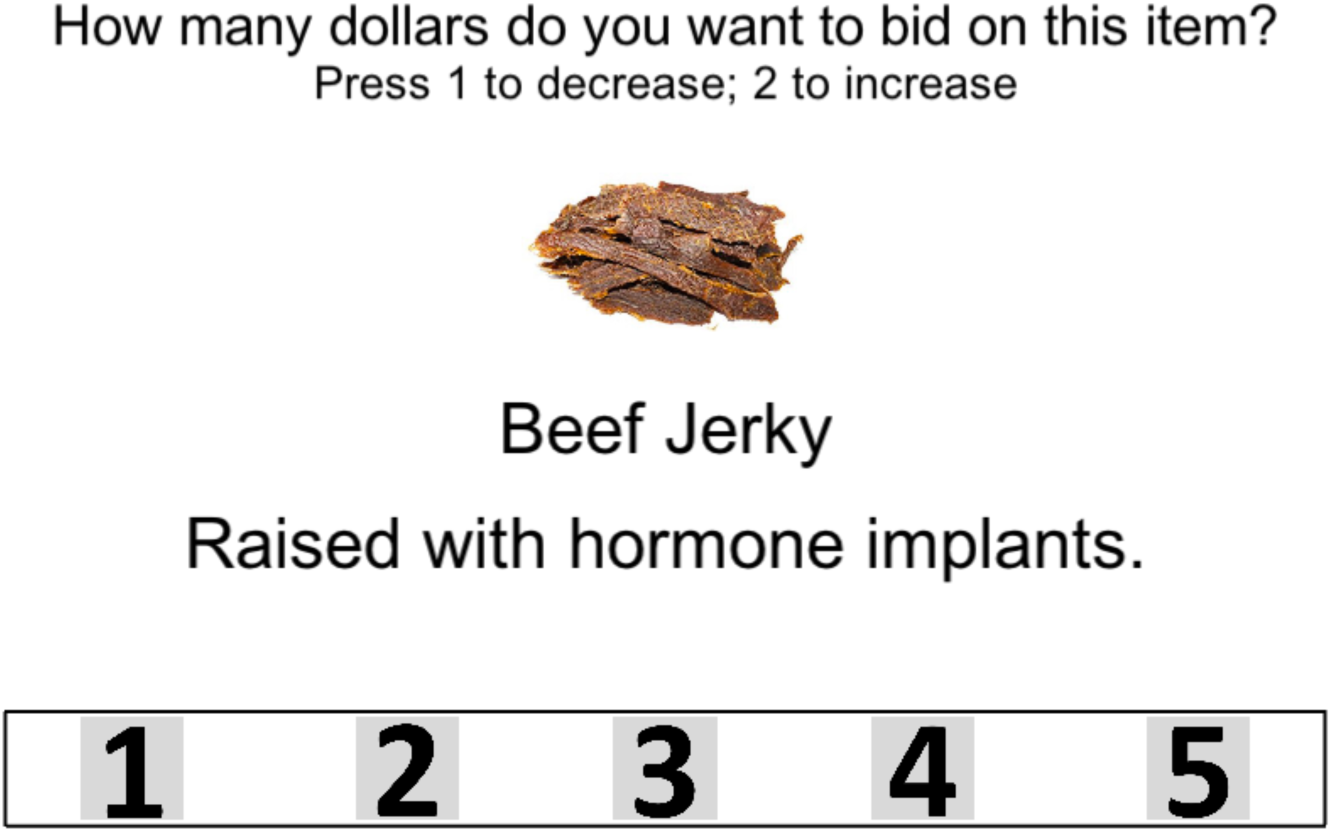
An example willingness-to-pay trial from the Becker-DeGroot-Marschack (1964) style auction in the neuroimaging study.

## References

1. Ajzen, I. (1991). The theory of planned behavior. Organizational behavior and human decision processes, 50(2), 179–211.

2. Badre, D., & D’esposito, M. (2009). Is the rostro-caudal axis of the frontal lobe hierarchical?. Nature Reviews Neuroscience, 10(9), 659–669.

3. Bak, H. J. (2001). Education and public attitudes toward science: Implications for the “deficit model” of education and support for science and technology. Social Science Quarterly, 82(4), 779–795.

4. Ballard, K., & Knutson, B. (2009). Dissociable neural representations of future reward magnitude and delay during temporal discounting. Neuroimage, 45(1), 143–150.

5. Bartra, O., McGuire, J. T., & Kable, J. W. (2013). The valuation system: a coordinate-based meta-analysis of BOLD fMRI experiments examining neural correlates of subjective value. Neuroimage, 76, 412–427.

6. Bechara, A., Damasio, H., & Damasio, A. R. (2000). Emotion, decision making and the orbitofrontal cortex. Cerebral cortex, 10(3), 295–307.

7. Becker, G. M., DeGroot, M. H., & Marschak, J. (1964). Measuring utility by a single-response sequential method. Behavioral science, 9(3), 226–232.

8. Bruce, A. S., Lusk, J. L., Crespi, J. M., Cherry, J. B. C., Bruce, J. M., McFadden, B. R., … & Martin, L. E. (2014). Consumers’ neural and behavioral responses to food technologies and price. Journal of Neuroscience, Psychology, and Economics, 7(3), 164–173

9. Christopoulos, G. I., Tobler, P. N., Bossaerts, P., Dolan, R. J., & Schultz, W. (2009). Neural correlates of value, risk, and risk aversion contributing to decision making under risk. Journal of Neuroscience, 29(40), 12574–12583.

10. Cox, D. N., & Evans, G. (2008). Construction and validation of a psychometric scale to measure consumers’ fears of novel food technologies: The food technology neophobia scale. Food quality and preference, 19(8), 704–710.

11. Crespi, J. M., Lusk, J. L., Cherry, J. B. C., Martin, L. E., McFadden, B. R., & Bruce, A. S. (2015). Neural activations associated with decision-time and choice in a milk labeling experiment. American Journal of Agricultural Economics, 98(1), 74–91.

12. Davis, T., Love, B. C., & Maddox, W. T. (2009). Anticipatory emotions in decision tasks: Covert markers of value or attentional processes?. Cognition, 112(1), 195–200.

13. Deichmann, R., Gottfried, J. A., Hutton, C., & Turner, R. (2003). Optimized EPI for fMRI studies of the orbitofrontal cortex. Neuroimage, 19(2), 430–441.

14. Delgado, M. R., Gillis, M. M., & Phelps, E. A. (2008). Regulating the expectation of reward via cognitive strategies. Nature neuroscience, 11(8), 880–881.

15. Feinstein, J. S., Adolphs, R., Damasio, A., & Tranel, D. (2011). The human amygdala and the induction and experience of fear. Current biology, 21(1), 34–38.

16. Frewer, L. J., Shepherd, R., & Sparks, P. (1994). Biotechnology and food production: knowledge and perceived risk. British Food Journal, 96(9), 26–32.

17. Friston, K. (2012). Ten ironic rules for non-statistical reviewers. Neuroimage, 61(4), 1300–1310.

18. Goel, V., & Dolan, R. J. (2003). Reciprocal neural response within lateral and ventral medial prefrontal cortex during hot and cold reasoning. Neuroimage, 20(4), 2314–2321.

19. Hart, W., Albarracín, D., Eagly, A. H., Brechan, I., Lindberg, M. J., & Merrill, L. (2009). Feeling validated versus being correct: A meta-analysis of selective exposure to information. Psychological Bulletin, 135(4), 555–588.

20. Hansen, J., Holm, L., Frewer, L., Robinson, P., & Sandøe, P. (2003). Beyond the knowledge deficit: recent research into lay and expert attitudes to food risks. Appetite, 41(2), 111–121.

21. Kable, J. W., & Glimcher, P. W. (2007). The neural correlates of subjective value during intertemporal choice. Nature neuroscience, 10(12), 1625–1633.

22. Kahathuduwa, C. N., Boyd, L. A., Davis, T., O’Boyle, M., & Binks, M. (2016). Brain regions involved in ingestive behavior and related psychological constructs in people undergoing calorie restriction. Appetite, 107, 348–361.

23. Kahathuduwa, C. N., Davis, T., O’Boyle, M., Boyd, L. A., Chin, S. H., Paniukov, D., & Binks, (2018). Effects of 3-week total meal replacement vs. typical food-based diet on human brain functional magnetic resonance imaging food-cue reactivity and functional connectivity in people with obesity. Appetite, 120, 431–441.

24. Kahnt, T., Heinzle, J., Park, S. Q., & Haynes, J. D. (2011). Decoding different roles for vmPFC and dlPFC in multi-attribute decision making. Neuroimage, 56(2), 709–715.

25. Kleijnen, M., Lee, N., & Wetzels, M. (2009). An exploration of consumer resistance to innovation and its antecedents. Journal of economic psychology, 30(3), 344–357.

26. Kober, H., Mende-Siedlecki, P., Kross, E. F., Weber, J., Mischel, W., Hart, C. L., & Ochsner, K. (2010). Prefrontal–striatal pathway underlies cognitive regulation of craving. Proceedings of the National Academy of Sciences, 107(33), 14811–14816.

27. Koechlin, E., Ody, C., & Kouneiher, F. (2003). The architecture of cognitive control in the human prefrontal cortex. Science, 302(5648), 1181–1185.

28. Krain, A. L., Wilson, A. M., Arbuckle, R., Castellanos, F. X., & Milham, M. P. (2006). Distinct neural mechanisms of risk and ambiguity: a meta-analysis of decision-making. Neuroimage, 32(1), 477–484.

29. Kraus, N., Malmfors, T., & Slovic, P. (1992). Intuitive toxicology: Expert and lay judgments of chemical risks. Risk analysis, 12(2), 215–232.

30. Kriegeskorte, N., Simmons, W. K., Bellgowan, P. S., & Baker, C. I. (2009). Circular analysis in systems neuroscience: the dangers of double dipping. Nature neuroscience, 12(5), 535–540.

31. Lang, J. T., & Hallman, W. K. (2005). Who does the public trust? The case of genetically modified food in the United States. Risk Analysis: An International Journal, 25(5), 1241–1252.

32. Levy, D. J., & Glimcher, P. W. (2012). The root of all value: a neural common currency for choice. Current opinion in neurobiology, 22(6), 1027–1038.

33. Linder, N. S., Uhl, G., Fliessbach, K., Trautner, P., Elger, C. E., & Weber, B. (2010). Organic labeling influences food valuation and choice. NeuroImage, 53(1), 215–220.

34. Lusk, J. L., Crespi, J. M., Cherry, J. B. C., Mcfadden, B. R., Martin, L. E., & Bruce, A. S. (2015). An fMRI investigation of consumer choice regarding controversial food technologies. Food Quality and Preference, 40, 209–220.

35. Lusk, J. L., Roosen, J., & Bieberstein, A. (2014). Consumer Acceptance of New Food Technologies: Causes and Roots of Controversies. Annual Review of Resource Economics, 6(1), 381–405.

36. McFadden, B. R., Lusk, J. L., Crespi, J. M., Cherry, J. B. C., Martin, L. E., Aupperle, R. L., & Bruce, A. S. (2015). Can Neural Activation in Dorsolateral Prefrontal Cortex Predict Responsiveness to Information? An Application to Egg Production Systems and Campaign Advertising. PloS one, 10(5), e0125243.

37. McClure, S. M., Laibson, D. I., Loewenstein, G., & Cohen, J. D. (2004). Separate neural systems value immediate and delayed monetary rewards. Science, 306(5695), 503–507.

38. Miller, E. K. (2000). The prefontral cortex and cognitive control. Nature reviews neuroscience, 1(1), 59–65.

39. Petty, R. E., Brinol, P. & Priester, J. R. (2009). Mass media attitude change: Implications of ” the elaboration likelihood model of persuasion. In J. Bryant & M. B. Oliver (Eds.), Media effects: Advances in theory and research (pp. 125–163). New York, NY: Routledge.

40. Petty, R. E., & Cacioppo, J. T. (1986). The elaboration likelihood model of persuasion. In Communication and persuasion (pp. 1–24). Springer, New York, NY.

41. Pinheiro, J., & Bates, D. (2006). Mixed-effects models in S and S-PLUS. Springer Science & Business Media.

42. Plassmann, H., O’Doherty, J., Shiv, B., & Rangel, A. (2008). Marketing actions can modulate neural representations of experienced pleasantness. Proceedings of the National Academy of Sciences, 105(3), 1050–1054.

43. Plassmann, H., & Weber, B. (2015). Individual differences in marketing placebo effects: Evidence from brain imaging and behavioral experiments. Journal of Marketing Research, 52(4), 493–510.

44. Platt, M. L., & Huettel, S. A. (2008). Risky business: the neuroeconomics of decision making under uncertainty. Nature neuroscience, 11(4), 398–403.

45. Pliner, P., & Hobden, K. (1992). Development of a scale to measure the trait of food neophobia in humans. Appetite, 19(2), 105–120.

46. Raichle, M. E. (2015). The brain’s default mode network. Annual review of neuroscience, 38, 433–447.

47. Rick, S. (2011). Losses, gains, and brains: Neuroeconomics can help to answer open questions about loss aversion. Journal of Consumer Psychology, 21(4), 453–463.

48. Roosen, J., Bieberstein, A., Blanchemanche, S., Goddard, E., Marette, S., & Vandermoere, F. (2015). Trust and willingness to pay for nanotechnology food. Food policy, 52, 75–83.

49. Roy, M., Shohamy, D., & Wager, T. D. (2012). Ventromedial prefrontal-subcortical systems and the generation of affective meaning. Trends in cognitive sciences, 16(3), 147–156.

50. Rushworth, M. F., & Behrens, T. E. (2008). Choice, uncertainty and value in prefrontal and cingulate cortex. Nature neuroscience, 11(4), 389–397.

51. Saba, A., Rosati, S., & Vassallo, M. (2000). Biotechnology in agriculture: perceived risks, benefits and attitudes in Italy. British Food Journal, 102(2), 114–122.

52. Schonberg, T., Fox, C. R., & Poldrack, R. A. (2011). Mind the gap: bridging economic and naturalistic risk-taking with cognitive neuroscience. Trends in cognitive sciences, 15(1), 11–19.

53. Sears, D. O., & Freedman, J. L. (1967). Selective exposure to information: A critical review. Public Opinion Quarterly, 31(2), 194–213.

54. Siegel, J. S., Power, J. D., Dubis, J. W., Vogel, A. C., Church, J. A., Schlaggar, B. L., & Petersen, S. E. (2014). Statistical improvements in functional magnetic resonance imaging analyses produced by censoring high-motion data points. Human brain mapping, 35(5), 1981–1996.

55. Siegrist, M. (2008). Factors influencing public acceptance of innovative food technologies and products. Trends in Food Science & Technology, 19(11), 603–608.

56. Siegrist, M., Cousin, M. E., Kastenholz, H., & Wiek, A. (2007). Public acceptance of nanotechnology foods and food packaging: The influence of affect and trust. Appetite, 49(2), 459–466.

57. Siegrist, M., Stampfli, N., Kastenholz, H., & Keller, C. (2008). Perceived risks and perceived benefits of different nanotechnology foods and nanotechnology food packaging. Appetite, 51(2), 283–290.

58. Slovic, P. (2016). The perception of risk. Routledge.

59. Smith, S. M., Fabrigar, L. R., & Norris, M. E. (2008). Reflecting on six decades of selective exposure research: Progress, challenges, and opportunities. Social and Personality Psychology Compass, 2(1), 464–493.

60. Sparks, P., Shepherd, R., & Frewer, L. J. (1995). Assessing and structuring attitudes toward the use of gene technology in food production: The role of perceived ethical obligation. Basic and applied social psychology, 16(3), 267–285.

61. Tenbült, P., de Vries, N. K., Dreezens, E., & Martijn, C. (2005). Perceived naturalness and acceptance of genetically modified food. Appetite, 45(1), 47–50.

62. Tenbült, P., de Vries, N. K., Dreezens, E., & Martijn, C. (2008). Intuitive and explicit reactions towards “new” food technologies: Attitude strength and familiarity. British Food Journal, 110(6), 622–635.

63. Tom, S. M., Fox, C. R., Trepel, C., & Poldrack, R. A. (2007). The neural basis of loss aversion in decision-making under risk. Science, 315(5811), 515–518.

64. Worthy, D. A., Davis, T., Gorlick, M. A., Cooper, J. A., Bakkour, A., Mumford, J. A., … & Maddox, W. T. (2016). Neural correlates of state-based decision-making in younger and older adults. NeuroImage, 130, 13–23.

